# Divergent Fates of Kidney-Resident Polyomaviruses: Stable Shedding Versus Near-Silent Persistence

**DOI:** 10.64898/2026.02.12.705499

**Authors:** Anik Mojumder, Kimin W. Nguyen, Christopher S. Sullivan

## Abstract

Polyomaviruses establish long-term infection in the kidney and are intermittently shed in urine. However, the relationship between kidney-resident viral genomes and urinary shedding during persistent infection remains poorly defined. Using a genetically barcoded murine polyomavirus library, we tracked thousands of viral lineages *in vivo* by pairing longitudinal urine sampling with endpoint barcode sequencing of kidney tissue in four mice. Across all animals, kidney infection consistently resolved into two stable viral populations, with near-silent persistence as the dominant fate. Most kidney-resident barcodes were never detected in late urine at late stages of infection, even though many reached substantial abundance within the kidney, demonstrating that kidney viral genome levels alone do not predict urinary shedding. In contrast, only a small minority of kidney barcodes contributed disproportionately to urine virus output at late timepoints, and these barcodes exhibited stable longitudinal behavior, with repeated detection in urine over time and markedly higher peak urine abundance than late non-shed or random barcode controls. Shedding behavior was not explained by input virus stock abundance, barcode sequence features, predicted miRNA targeting, or ongoing reseeding from blood or other tissues. Instead, barcodes that ultimately dominated late urine already showed elevated urine detection early after infection, indicating that shedding fate is established early and maintained throughout persistent infection. Together, these findings reveal that persistent kidney infection is a structured reservoir composed of a large population of deeply restricted viral genomes and a smaller, stable subset that repeatedly produces urine-detectable viruses, with concurrent smoldering infections and latency-like restriction representing one possible model to explain the sharply different probabilities of shedding among kidney-resident genomes.

## Introduction

Polyomaviruses (PyVs) are among the best studied tumor viruses and have served for decades as foundational model systems for understanding viral oncogenesis, DNA replication, and host-virus interactions [1–7]. Consistent with their evolutionary success, 50-90% of adults are seropositive for one or more PyVs, reflecting widespread lifelong exposure [8]. Despite this, a fundamental aspect of PyV biology remains poorly understood: how do these simple viruses, with DNA genomes of only ∼5 kb, persist within their hosts for decades? Clinically, this question carries major consequences. PyV persistence and reactivation underlie severe disease in immunocompromised settings, including BK polyomavirus (BKPyV)-associated nephropathy and graft failure following kidney transplantation, as well as JC polyomavirus (JCPyV)-driven progressive multifocal leukoencephalopathy [9–13]. Although Merkel cell polyomavirus (MCPyV) is the only human PyV definitively established as a direct cause of cancer, through its role in Merkel cell carcinoma of the skin, emerging evidence suggests that persistent BKPyV infection contributes to bladder carcinogenesis [14–20]. Therefore, from both clinical and basic biology perspectives, there is a critical need to understand how PyVs establish and maintain long-term infection *in vivo*.

Urinary shedding is a defining feature of kidney-tropic PyVs and a clinically relevant non-invasive readout of viral replication activity during persistent infection [21–23]. In humans, BKPyV and JCPyV preferentially persist in the kidney and are shed into the urine, where longitudinal studies reveal typically episodic high-level shedding interspersed with periods of minimal or undetectable viral output [24–27]. In this context, because it recapitulates kidney tropism, long-term persistence, and urinary shedding in a tractable experimental setting, murine polyomavirus (muPyV) provides a powerful *in vivo* system to interrogate the mechanisms underlying these shedding patterns [28–31].

With the advent of next-generation sequencing technologies, we developed a genetically barcoded murine polyomavirus (muPyV) library that preserves viral fitness and enables longitudinal non-invasive simultaneous tracking of thousands of distinct viral lineages *in vivo* [31,32]. Using this barcoded muPyV system in our previous work, we discovered at least two distinct modes of urinary shedding during persistent infection: a continuous, low-level shedding (“smoldering”) involving initially thousands and then hundreds of viral barcodes and episodic high-level shedding events dominated by one or a few barcodes at later times of persistent infection [31]. However, our original study did not answer several fundamental questions: During long-term infection, which viral genomes persisting in the kidney ultimately give rise to urinary shedding? Do most kidney-resident viral genomes contribute to urine virus loads, or do many persist without shedding? Among those that do shed, is long-term persistent shedding driven by a stable subset of kidney-resident viral lineages or by transient contributions from different viral genomes over time? Does kidney viral abundance predict long-term shedding fate? These open questions formed the basis of the present study.

Current understanding of PyV shedding patterns exhibits hallmarks consistent with potential latency and reactivation [24]. PyV persistence has long been connected to immune control [29,33–36]. Only recently has PyV persistence been discussed in the context of basic host cell-cycle-mediated regulation [24,37–42]. Although classical models proposed that large T antigen (LTAg) actively drives infected cells into S phase, recent work on BKPyV instead demonstrates that entry into S phase is required for robust LTAg expression [24,37,38,40,42–44]. Consistent with this model, BKPyV genomes remain largely transcriptionally quiescent in G_0_-arrested cells, while progression from G_₁_ into S phase coincides with activation of early viral gene expression, possibly in part through release of E2F transcription factors from RB and their engagement with binding sites in the viral non-coding control region (NCCR) [24,37,38]. Once activated, BKPyV may further reinforce a replication-competent state by engaging DNA-damage response pathways and reinforcing derepressed “free” E2F that prolong S phase [39,45,46].

In parallel, host immune signaling further shapes persistent infection. BKPyV infection induces strong IRF3-driven interferon responses in endothelial cells but not in renal epithelial cells, creating a cellular environment permissive for long-term persistence in the kidney [40]. Single-cell transcriptomic studies reinforce this view, revealing a consistent host signature of productive infection characterized by S/G2M cell-cycle programs, DNA-damage and replication-stress pathways, MAPK activation, mitochondrial stress, and suppression of interferon-stimulated and antigen-presentation genes [41,47–52]. Importantly, only a minority of infected cells become high producers with robust late capsid transcription, while the majority remain in low or restricted transcriptional states despite similar viral exposure [49]. Together, these observations led us to hypothesize a subset of virus lineage reservoirs are primed for continuous/frequent shedding with other viral genomes persisting in the kidney in deeply restricted states that infrequently reactivate contributing to the urine virome.

Here, we tested this hypothesis using our longitudinal *in vivo* muPyV barcode sequencing data from four individual mice (FL, FR, ML, and MR) [31] applying new analyses to classify kidney-resident viral genomes according to their long-term shedding behavior. By partitioning kidney barcodes into shedding and non-shedding populations at latest times of infection within each mouse and tracking their dynamics over time from initial infection, we determined whether shedding fate is associated with kidney abundance, barcode sequence features, or ongoing reseeding from other tissues, and assessed when during infection these fates emerge within individual hosts. Across all four mice, we find that persistent kidney infection is consistently structured into two stable viral populations: a minority of genomes that drive sustained and episodic urinary shedding and a majority that persist without contributing to long-term bulk viral genome urine output. These findings demonstrate a large population of kidney resident PyV lineages that rarely or never give rise to shed viruses and that shedding fate is established early during infection and maintained over time during long-term persistent infection *in vivo*.

## Materials and Methods

### Generation of barcoded virus and longitudinal infection model

We previously generated a complex library of barcoded muPyV in which each viral genome carries a unique 12-nucleotide random barcode inserted at a neutral position between the two polyadenylation signals [31,32]. This design allows individual viral barcodes to be tracked across tissues and over time without perturbing viral replication or fitness. Barcoded muPyV stocks were generated by excising viral genomes from plasmids, re-circularizing them, and propagating virus in NMuMG cells, described previously [31,32].

All animal procedures were previously published [31]. Animal procedures were conducted in accordance with protocols approved by The University of Texas at Austin Institutional Animal Care and Use Committee (AUP-2022–00305) and were subject to veterinary approval. To study longitudinal shedding dynamics and organ-resident viral populations, we infected two female (“FL” and “FR”) and two male (“ML” and “MR”) FVB/NJ mice (9-10 weeks old) with 1 × 10:J infectious units of the barcoded muPyV library via intraperitoneal injection. We collected urine non-invasively at multiple timepoints over the course of infection and stored samples for downstream barcode sequencing and viral load quantification. At the end of the study (59 days post-infection for mouse “MR” and 99 days post-infection for the remaining three mice), we harvested a broad panel of organs to capture the full tissue distribution of viral barcodes. Mouse “MR” was euthanized earlier due to the development of an abscess that was determined to be unrelated to muPyV infection. We purified DNA from urine, tissues, and blood and quantified total viral genome abundance in each sample by real-time qPCR. Barcode identities and relative abundances were determined by targeted amplification of the barcode-containing region followed by Illumina sequencing. Briefly, barcode amplicon libraries were generated using enrichment and indexing PCRs and sequenced on an Illumina NextSeq platform. All experimental procedures have been described in detail in our prior work [31].

### Extracting barcode sequences

We extracted barcode sequences from FASTQ files as previously described [31]. Briefly, we used Cutadapt to identify barcodes, allowing the default error rate of 10% and requiring both flanking adapters to be present via linked-adapter matching [53]. We then aggregated raw barcode counts for each sample using custom R scripts built on tidyverse packages [54,55].

### Clustering and defining the stock barcode reference

To define a reference set of input barcodes, we clustered barcodes from the stock virus using Starcode message-passing clustering, with a maximum Levenshtein distance of 3 and the default cluster ratio of 5, as in our prior work [31,32,56]. We then restricted downstream analyses to the subset of barcodes accounting for the top 99% of cumulative abundance in the stock, excluding low-complexity artifacts. This yielded an estimated 4012 unique barcodes in the initial barcoded virus stock, which served as the reference set for all barcode matching and quantification **(Table S1)**. Unless otherwise noted, we performed all downstream processing and analyses in R (version 4.3.3) using tidyverse [54] and associated packages (including patchwork [57], ggridges [58], ComplexUpset [59], pheatmap [60], corrplot [61], kableExtra [62], stringdist [63], purrr [64], reshape2 [65], and gridExtra [66]).

### Assigning sample barcodes to stock barcodes

To quantify barcode abundance in urine and tissue samples, we mapped every observed sample barcode to the closest stock barcode based on Levenshtein distance. We discarded sample barcodes that did not fall within a distance of 3 from any stock barcode. When a sample barcode matched multiple stock barcodes at the same minimum distance, we divided its count evenly among those stock barcodes. We then summed these fractional assignments within each sample to obtain a weighted count for each stock barcode, reflecting its estimated contribution to that biological sample. To reduce the influence of extremely low-abundance signals, we removed weighted barcode counts below 10. This cutoff served as our detection threshold for subsequent barcode-level analyses.

### Normalizing relative barcode abundance to viral genome load

Because total viral genome abundance varied across biological samples, we normalized barcode counts to independent qPCR-based measurements of viral load (approximated by viral genome amounts detected). For each sample, we first calculated the fraction of total barcode signal (defined as weighted barcode counts after barcode-to-stock assignment) contributed by each stock barcode. We then scaled this fraction by the measured viral genome concentration in that sample (barcode fractions x genome level) to obtain an estimated barcode-specific viral load. We refer to this scaled quantity as the “barcode level”, which represents the estimated abundance of an individual viral barcode after accounting for both its relative representation within a sample and the absolute viral genome load of that sample. For urine samples, we multiplied each barcode’s fractional abundance by the qPCR-estimated number of viral genomes per microliter of urine, yielding barcode-specific genome concentrations in urine **(Table S2)**. For tissue samples, we multiplied fractional barcode abundance by qPCR-estimated viral genomes per microgram of input DNA **(Table S3)**. For blood samples, we scaled fractional barcode abundance by viral genome concentration per microliter of blood and converted values to a per-milliliter basis to maintain consistency across sample types **(Table S3)**.

### Definition of late urine detection and kidney barcode classification

For each animal, we defined the late window as the final two urine collection timepoints prior to sacrifice (FL: days 90 and 98; FR: days 94 and 98; ML: days 94 and 98; MR: days 55 and 59 post-infection). We classified a kidney barcode as “late-shed” if detectable in urine at either of these timepoints and as “late non-shed” if it was not selected during this window. We identified kidney-resident barcodes from tissue sequencing data by selecting barcodes detected in kidney samples. We assigned late-shed and late non-shed status independently for each mouse based on detection in urine.

### Quantification of kidney barcode counts, total load, and per-barcode abundance

For each mouse, we quantified the number of distinct kidney barcodes classified as late-shed or late non-shed (see “Definition of late urine detection and kidney barcode classification”). To quantify the total kidney barcode load for each group, we summed the barcode-level (scaled quantity estimate of total barcode genome abundance) values across all barcodes belonging to the late-shed group and separately across all barcodes belonging to the late non-shed group within each mouse. Note that each barcode was associated with a barcode level, defined as the qPCR-scaled estimate of viral genome abundance for that barcode (see “Normalizing barcode abundance to viral genome load”), expressed as viral genomes per microgram of input kidney DNA. To test whether kidney barcode abundance alone predicted late urinary shedding, we compared the distributions of kidney barcode levels per-barcode between late-shed and late non-shed groups within each mouse. Here, per-barcode kidney barcode level refers to the individual barcode-level values (qPCR-scaled estimate of viral genomes per microgram of kidney DNA) for each barcode, analyzed as distributions of kidney barcode levels per-barcode rather than summed totals. This analysis directly tests whether individual viral genomes that contribute to late urine shedding are distinguished simply by higher abundance in the kidney.

### Urine barcode composition and contribution to total urine viral load

We analyzed longitudinal urine barcode composition using two complementary approaches. First, for each urine sample, we calculated barcode fractional abundance, defined as the weighted barcode count for a given barcode divided by the sum of the weighted barcode counts across all barcodes detected in that urine sample (fractional abundance for barcode *i* = weighted barcode count for barcode *i* / total weighted barcode counts in that sample). Note, weighted barcode counts reflect post-clustering, tie-resolved barcode counts after removal of low-abundance signals (see “Assigning sample barcodes to stock barcodes”). Then, the fractional abundances were normalized within each urine sample and were used to generate donut plots visualizing barcode composition at each timepoint. In these plots, we highlighted the top 50 kidney barcodes, defined as the 50 barcodes with the highest kidney barcode levels within the specified group (late-shed or late non-shed) for each individual mouse. These highlighted barcodes were assigned unique colors, while all remaining barcodes detected in urine were displayed in alternating grayscale to preserve rank ordering while minimizing overplotting. Second, to quantify how much of the urine signal was contributed by these kidney barcode sets, we performed a urine contribution analysis, in which we summed urine barcode levels across the selected barcodes at each timepoint. Urine barcode level refers to the qPCR-scaled estimate of barcode abundance in urine, expressed as viral genomes per microliter of urine (see “Normalizing relative barcode abundance to viral genome load”). For each timepoint, we summed urine barcode levels for the top 50 kidney barcodes and overlaid this value on the total urine viral signal, defined as the sum of urine barcode levels across all barcodes detected in that urine sample.

### Barcode-resolved longitudinal urine trajectory analysis

To determine whether late urinary shedding reflected consistent, barcode-specific shedding behavior over time (i.e., repeated detection of the same viral lineages across multiple urine collections, rather than transient or sporadic appearance) behavior, we analyzed longitudinal urine trajectories at the level of individual viral barcodes tracked independently over time. We examined three barcode groups. These included (i) the top 50 most abundant late-shed kidney barcodes, defined as the 50 kidney-resident barcodes with the highest kidney barcode levels (qPCR-scaled estimate of viral genomes per microgram of kidney DNA) among barcodes classified as late-shed, (ii) the top 50 late non-shed kidney barcodes defined as the 50 kidney-resident barcodes with the highest kidney barcode levels among barcodes classified as late non-shed, and (iii) a random barcode control group of equal size per mouse, drawn from the set of all barcodes detected in urine after excluding barcodes belonging to the first two groups. For each barcode, we quantified its longitudinal urine shedding trajectory by tracking its urine barcode levels (qPCR -scaled estimate of viral genomes per microliter of urine) at each urine collection timepoint. For visualization only, we capped barcode level values at an upper limit of 25 to improve interpretation of the trajectories.

### Input virus stock relative barcode rank analysis

To assess whether late shedding behavior reflected differences in input abundance, we ranked viral barcodes by their abundance in the input virus stock, defined as the weighted barcode count obtained from sequencing of the barcoded virus stock prior to infection. We derived stock ranks from the set of barcodes comprising the top 99% of cumulative abundance in the input library, ordering barcodes from most abundant (rank 1) to least abundant. For each mouse, we examined stock rank distributions for the top 50 late-shed and top 50 late non-shed kidney barcodes , where “top 50” refers to the 50 barcodes within each group with the highest kidney barcode levels (qPCR-scaled estimate of viral genomes per microgram of kidney DNA) in that mouse and summarized central tendencies by calculating median stock ranks for each group.

### Peak urine barcode abundance analysis

To identify barcodes associated with high-amplitude urinary shedding, we calculated the peak urine abundance for each barcode, defined as the maximum urine barcode level observed for that barcode across all urine collection timepoints. Here, urine barcode level refers to the estimated abundance of a barcode in urine after normalizing its sequencing representation to the total viral genome concentration measured by qPCR, expressed as viral genomes per microliter of urine (see “Normalizing barcode abundance to viral genome load”). We performed this analysis for the top 50 late-shed kidney barcodes, the top 50 late non-shed kidney barcodes, and a size-matched random barcode control group, with all groups defined independently for each mouse. Here, “top 50” refers to the 50 barcodes within each group with the highest kidney barcode levels (qPCR-scaled estimate of viral genomes per microgram of kidney DNA). We log_₁₀_-transformed peak urine abundance values for visualization. We performed comparisons within individual mice, such that peak urine abundance distributions for different barcode groups were compared only among barcodes originating from the same animal, thereby accounting for mouse-to-mouse differences in overall viral shedding levels.

### Definition and enrichment analysis of smoldering barcodes

To identify barcodes exhibiting sustained shedding (“smoldering”), we quantified barcode detection frequency across urine timepoints, defined as the proportion of urine collection timepoints at which a given barcode was detected above the barcode-level detection threshold. Detection was defined as a urine barcode level corresponding to a weighted barcode count ≥ 10 after barcode-to-stock assignment, consistent with thresholds used throughout the study (see “Assigning sample barcodes to stock barcodes”). For each barcode within a given mouse, we calculated detection frequency as the number of urine timepoints with detection divided by the total number of urine timepoints sampled for that mouse. Within each mouse, we defined the top 50 barcodes with the highest detection frequency as “top smolderers”. We compared the fraction of top smoldering barcodes belonging to the late-shed or late non-shed kidney barcode groups against a size-matched random barcode control.

### Early versus late urine detection frequency analysis

To test whether barcodes that dominate late urine already exhibit elevated activity earlier in infection, we partitioned urine time courses into early and late phases. For each mouse, we defined the late phase as the final quartile of urine sampling timepoints closest to the time of sacrifice (e.g., ≥ day 74 for mice FL, FR, and ML; ≥ day 45 for mouse MR) , with all earlier timepoints assigned to the early phase. We then grouped barcodes into three sets. These included top late smolderers, defined as the 50 barcodes with the highest detection frequency during the late phase (detection frequency defined as the number of urine timepoints at which a barcode was detected above the barcode-level detection threshold divided by the total number of late-phase urine timepoints for that mouse); top late high-shedders, defined as the 50 barcodes with the greatest cumulative qPCR-scaled urine barcode levels (viral genomes per microliter of urine) across all late-phase urine timepoints and a size-matched random barcode control group, defined independently for each mouse. For each barcode, we calculated the fraction of early-phase detection frequency (defined as the number of early-phase urine timepoints at which a barcode was detected above the barcode-level detection threshold divided by the total number of early-phase urine timepoints for that mouse).

### Correlation analysis between kidney barcode groups and other tissues

To examine relationships between kidney barcode populations and other tissues, we computed Spearman correlation matrices using barcode relative abundance values, defined as the fractional abundance of each barcode within a given tissue sample (weighted barcode count for a barcode divided by the sum of weighted barcode counts across all barcodes in that tissue sample). For this analysis, we split kidney barcodes into late-shed and late non-shed groups using the same classification criteria described earlier, where late-shed kidney barcodes were those detected in urine during the final two urine timepoints for a given mouse, and late non-shed kidney barcodes were those not detected during this window (see “Definition of late urine detection and kidney barcode classification”). We treated these two groups as separate pseudo-organs. We computed Spearman correlations using pairwise complete observations and visualized correlation matrices using mixed circle-and-number correlograms. We also examined Spearman correlations across mice using “kidney-only” barcode data, defined as comparisons of relative barcode abundance profiles in kidney tissue between different mice, restricted to barcodes detected in kidney samples. This analysis tested whether the composition of late-shed or late non-shed kidney barcode populations was conserved across animals.

### Barcode sequence feature analyses

To test whether intrinsic barcode sequence features contributed to shedding fate, we analyzed GC content, barcode length, and predicted miRNA targeting. For each mouse, we computed GC content and barcode length distributions for the top 50 most abundant late-shed and top 50 late non-shed kidney barcodes, where “top 50” refers to the 50 barcodes within each group with the highest kidney barcode levels (qPCR-scaled viral genome abundance in kidney tissue) for that mouse. To assess potential miRNA-mediated effects, we tested whether kidney barcodes were enriched for predicted miRNA seed matches. We defined a miRNA-hit barcode as any barcode whose sequence contained a canonical miRNA seed match, defined as perfect Watson-Crick complementarity to nucleotides 2-8 from the 5′ end of a mature miRNA. This region corresponds to the miRNA seed sequence, which is widely recognized as a primary determinant of miRNA target recognition and binding specificity in animals [67–69]. For each miRNA, we searched for perfect matches to nucleotides 2-8 within both the forward barcode sequence and a corresponding reverse-orientation barcode sequence generated for each barcode using custom Python scripts. Seed matches identified in the reverse orientation were collapsed back to their corresponding forward barcode identity. We performed this analysis separately for host murine miRNAs curated in miRbase [70] and for the muPyV-encoded miRNAs described previously [71], restricting the murine analysis to the top 100 kidney-abundant miRNAs extracted from the published small RNA sequencing atlas [72]. We compiled the set of miRNA-hit barcodes by retaining unique forward barcode sequences associated with at least one seed match and intersected this set with kidney-resident barcodes for each mouse. For each mouse, miRNA targeting was quantified using two complementary metrics: an unweighted metric corresponding to the fraction of kidney barcodes classified as late-shed or late non-shed that contained at least one miRNA seed match, and an abundance-weighted metric corresponding to the fraction of total kidney viral genome abundance (barcode level) attributable to miRNA-hit barcodes within each group. To determine whether observed differences in miRNA targeting between late-shed and late non-shed groups could be explained by intrinsic barcode sequence properties, we generated matched random expectations independently for each mouse using a length- and GC-matched resampling strategy. Within each mouse, kidney barcodes were stratified by exact barcode length and by GC content, with GC content divided into five quantile-based bins to ensure approximately equal numbers of barcodes per bin. For each shedding group, the observed barcode set was characterized by its joint length and GC-bin composition, and a matched random set of equal size was generated by sampling barcodes from the full kidney barcode pool of the same mouse while preserving the exact counts within each length × GC-bin stratum.

Unweighted and abundance-weighted miRNA targeting metrics were recalculated for each resampled set, and this procedure was repeated 2,000 times per mouse and shedding group to generate empirical null distributions representing the expected range of miRNA targeting under random assignment of shedding fate while preserving barcode sequence composition.

All custom scripts used in this study are publicly available at https://github.com/ChrisSullivanLab/Shedding-dynamics-of-a-DNA-virus. Independent analyses validating trends of plots presented in this work, including confirmatory plots and accompanying documentation generated by one of the authors (K.W.N.), are included in the same repository. Tables of analyses of raw sequencing making up the plots in this work are provided in the Supplementary Tables file.

## Results

### Late urinary shedding arises from a minority of kidney barcodes, revealing a large non-shedding reservoir

To determine how kidney-resident viral barcodes contribute to late urinary output, we classified kidney barcodes based on their detection during the final phase of infection. Because kidneys were harvested at the experimental endpoint, we focused on the final two urine timepoints as the most proximal readout of kidney-derived viral shedding. Kidney barcodes detected in either of these timepoints were classified as “late-shed”, whereas kidney barcodes never detected during this window were classified as “late non-shed”.

We found that late non-shed barcodes greatly outnumbered late-shed barcodes across all four mice (**Fig. 1A; Table S4**). In FL, we identified 1,305 late non-shed distinct kidney barcodes compared with 188 distinct late-shed barcodes. We observed similar patterns in FR (913 late non-shed vs. 516 late-shed), ML (1,333 vs. 241), and MR (1,087 vs. 78) **(Table S4)**. Thus, in every mouse, late urinary shedding arose from only a minority of kidney barcode diversity, indicating that most viral genomes persisting in the kidney do not contribute detectably to late urine.

**Fig. 1.**
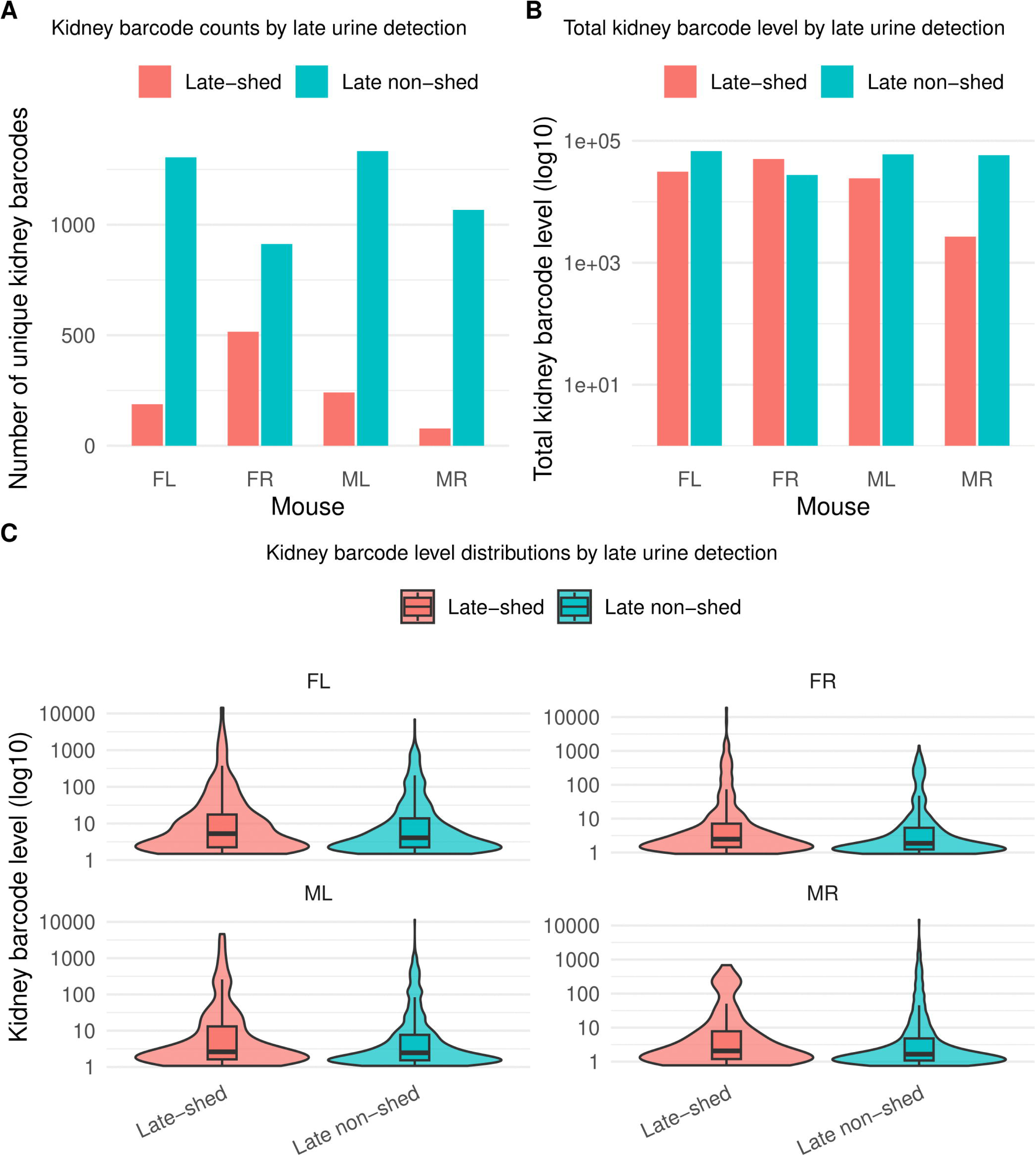
Late urinary shedding derives from a minority of kidney-resident viral barcodes. Kidney viral barcodes were classified based on their detection in urine during the final phase of infection, with kidney barcodes detected in either of the final two urine timepoints defined as **late-shed** and kidney barcodes never detected during this window defined as **late non-shed**. Shown in **(A)** is the number of unique kidney barcodes per mouse (FL, FR, ML, MR) stratified by late urine detection status. Shown in **(B)** is the total kidney barcode level for late-shed and late non-shed groups in each mouse, calculated as the summed barcode read counts and plotted on a log_₁₀_ scale. Shown in **(C)** are the distributions of per-barcode kidney abundance (log_₁₀_ scale) for late-shed and late non-shed barcodes within each mouse, with box plots indicating the median and interquartile range. Together, these data show that late urinary shedding arises from a small subset of kidney-resident viral genomes and that kidney barcode abundance alone does not distinguish barcodes that do versus do not contribute to late urine.

We next asked whether late non-shed barcodes represent rare kidney residents or instead form a major quantitative reservoir of the virus genomes (bulk total viral DNA) within the kidney. For each mouse, we summed kidney barcode levels separately for late-shed and late non-shed barcode groups (**Fig. 1B; Table S5**). Barcode levels are expressed as qPCR-scaled viral genome abundance per microgram of kidney DNA for each barcode (see Materials and Methods). In three of four mice (FL, ML, and MR), late non-shed barcodes contributed the majority of total kidney viral genome abundance **(Table S5).** FR differed, with higher total kidney barcode levels among late-shed barcodes than late non-shed barcodes **(Table S5)**. Importantly, even in FR, late-shed barcodes comprised fewer distinct kidney barcodes than late non-shed barcodes, indicating that absence from late urine is not simply a consequence of low abundance in the kidney.

We further tested whether kidney barcode abundance alone predicts late urinary shedding. For each barcode, we quantified its abundance by calculating its barcode level within the kidney of a given mouse, yielding an estimate of the total kidney-associated viral genome load contributed by that individual barcode. We then compared these per-barcode kidney abundance values between late-shed and late non-shed groups within each mouse (**Fig. 1C; Table S6**). Across all animals, we observed wide dynamic ranges and substantial overlap between groups. While some high-abundance kidney barcodes were late-shed, many barcodes with comparable kidney levels remained late non-shed. Thus, kidney barcode abundance alone is insufficient to explain whether a viral genome contributes to late urinary shedding.

### Late-shed but not late non-shed kidney barcodes contribute substantially to persistent urinary shedding

In our previous work, population-level analyses showed that although many viral genomes are detectable in urine, shedding during long-term persistent infection is dominated by a small subset of barcodes [31]. Here, we asked whether this principle persists when kidney-resident viral genomes are stratified by late shedding behavior. To address this, we examined longitudinal urine shedding patterns for kidney barcodes classified as late-shed or late non-shed, focusing on the most abundant kidney barcodes within each group (top 50 per mouse). For each urine sample, we calculated relative abundance of each barcode, defined as the fraction of total barcode-derived abundance within that urine sample (i.e., barcode-specific weighted counts normalized per timepoint). These relative compositions were visualized using donut plots (**Fig. 2A–B**). In parallel, we quantified absolute contributions to urine shedding by summing barcode levels and visualized these values over time using sum-overlay plots (**Fig. 2C–D**).

**Fig. 2.**
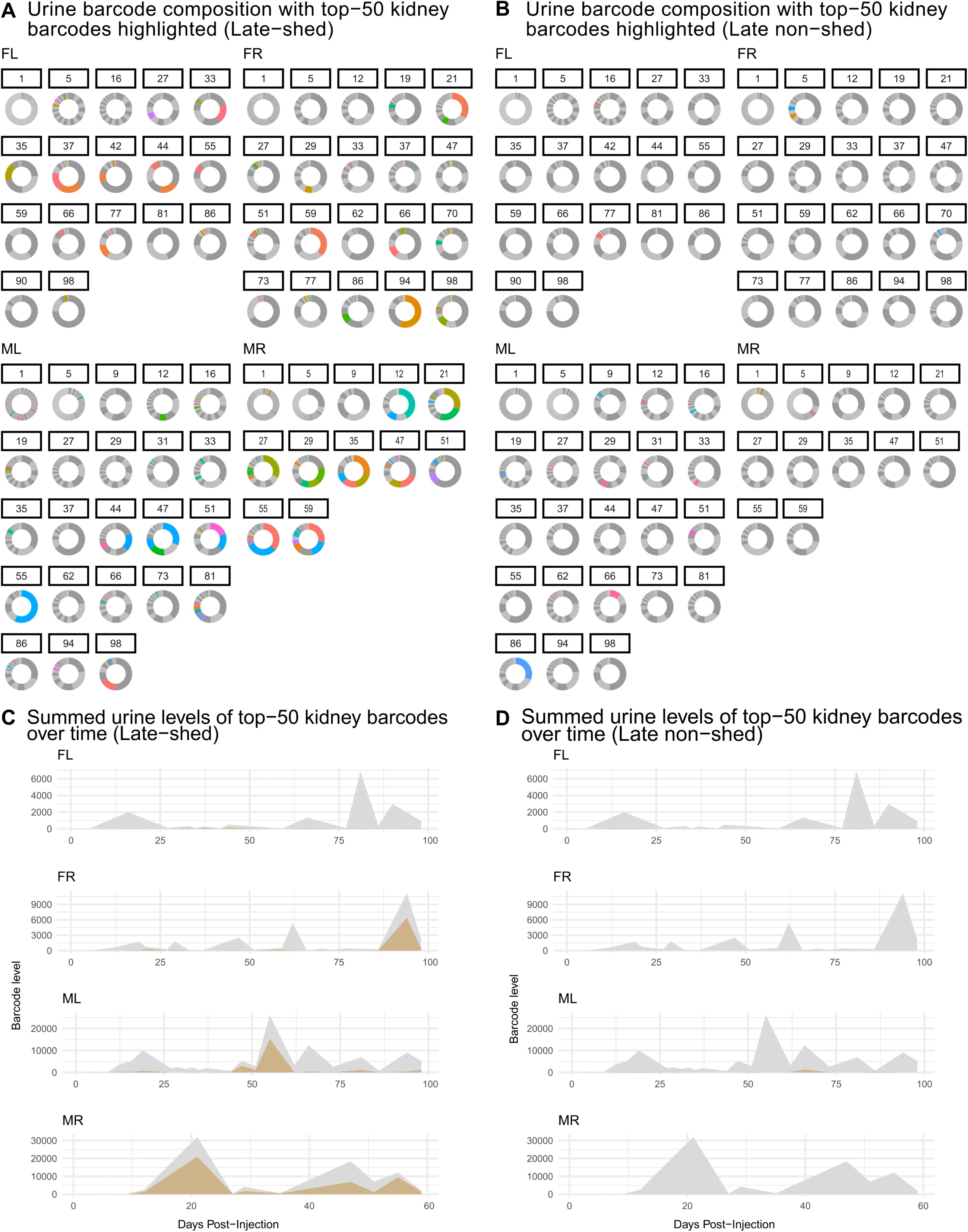
Longitudinal urine barcode composition for late-shed and late non-shed kidney barcodes. Kidney barcodes were classified as late-shed or late non-shed based on their detection in the final two urine timepoints. Shown in **(A)** and **(B)** are donut plots depicting urine barcode composition over time for late-shed and late non-shed kidney barcodes, respectively. For each urine timepoint, individual barcodes are represented as segments of a donut, with the area of each segment proportional to that barcode’s abundance relative to the total urine barcode DNA present at that timepoint. Most barcodes are present at low relative abundance and appear as dark or light gray segments that collectively form a largely uniform gray background. Colored segments represent individual barcodes that were among the top 50 most abundant kidney barcodes within the indicated group for each mouse. Shown in **(C)** are sum-overlay plots showing, for each mouse, the summed urine barcode levels over time of the top 50 late-shed kidney barcodes (gold), overlaid on the total bulk urine barcode levels at each timepoint (gray). Shown in **(D)** are analogous sum-overlay plots for the top 50 late non-shed kidney barcodes. These plots show that longitudinal urine shedding is largely accounted for by late-shed kidney barcodes, whereas late non-shed kidney barcodes remain weakly represented.

We observed that urine shedding during the persistent phase of infection (post-day 27) increasingly reflected contributions from a restricted number of late-shed kidney barcodes across all four mice (**Fig. 2A–B; Table S7-8**). These barcodes appeared as colored segments corresponding to the top 50 most abundant barcodes. In contrast, donut plots for late non-shed kidney barcodes showed little to no contribution from colored barcodes at late timepoints, reflecting their consistently low fractional representation in urine. Thus, while many barcodes remained detectable, only late-shed kidney barcodes rose to high fractional abundance within the urine barcode population throughout persistent infection.

Using sum-overlay analyses, we found that late-shed kidney barcodes contributed a substantially larger share of total urine barcode levels during the persistent phase (post-day 27) of infection compared to late non-shed kidney barcodes in all four mice (**Fig. 2C–D; Table S9-10**). Late non-shed kidney barcodes contributed minimally to cumulative urine shedding over time, even when they were abundant in kidney, indicating that kidney residence alone does not predict contribution to sustained urinary shedding.

### Late-shed barcodes are longitudinally consistent throughout persistent infection

Population-level dominance of late-shed kidney barcodes raised the question of whether this pattern reflects consistent longitudinal behavior of individual viral lineages or transient effects at late times of persistence. To address this, we analyzed urine shedding trajectories at per-barcode resolution for kidney late-shed barcodes, kidney late non-shed barcodes, and a random barcode control group matched in number (top 50 per mouse; **Fig. 3; Table S11-12**). We visualized longitudinal urine detection for individual barcodes across all timepoints to assess persistence and recurrence of shedding behavior.

**Fig. 3.**
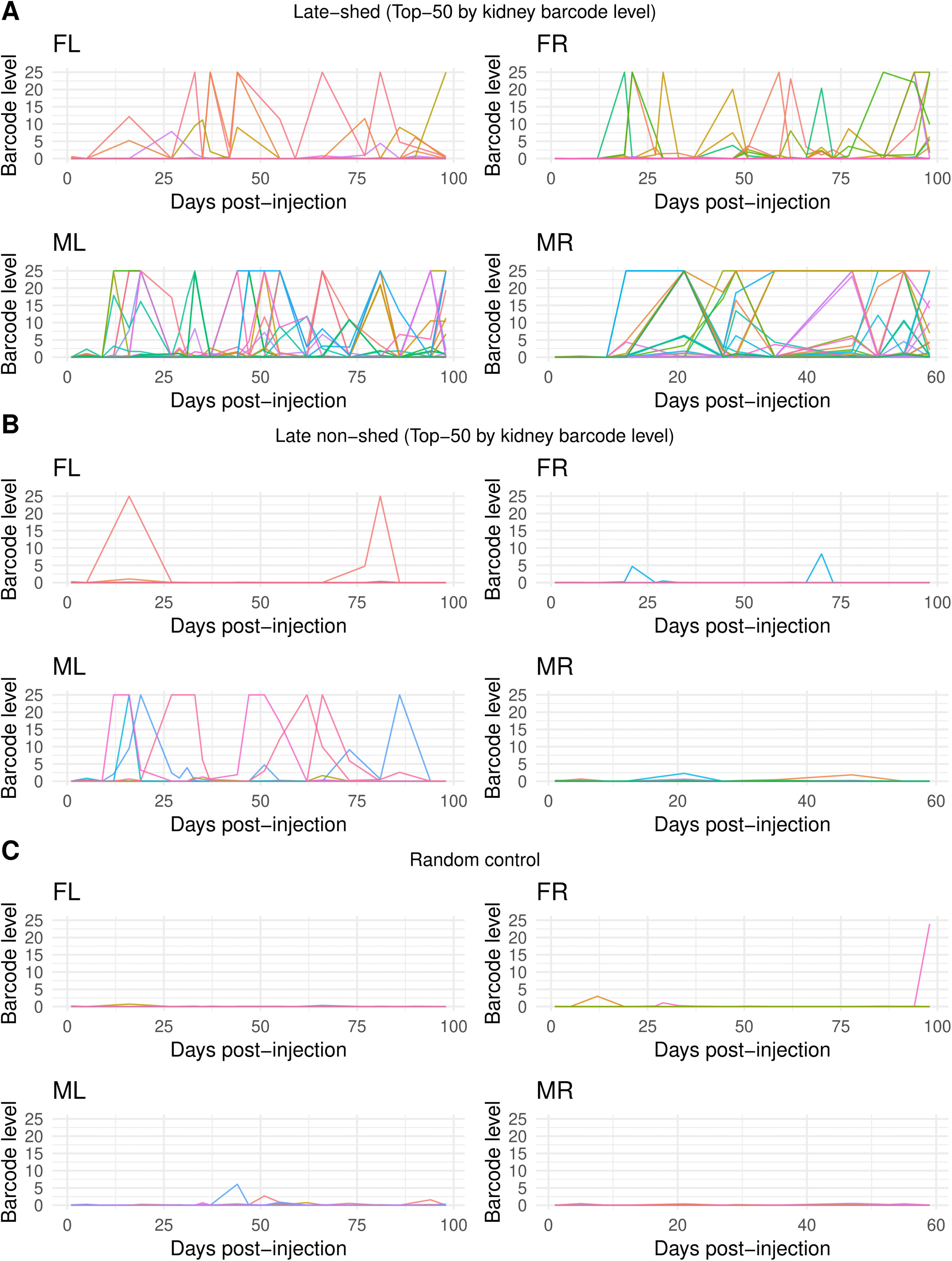
Barcode-resolved longitudinal urine shedding patterns for late-shed, late non-shed, and random kidney barcodes. Shown are barcode-resolved longitudinal urine shedding trajectories plotted over time for individual barcodes in each mouse. In **(A)**, lines represent urine barcode levels for the top 50 late-shed kidney barcodes per mouse, with each colored line corresponding to a single barcode tracked across all urine collection timepoints. In **(B)**, analogous plots are shown for the top 50 late non-shed kidney barcodes per mouse. In **(C)**, plots are shown for a size-matched random control group of 50 barcodes per mouse. For all panels, the x-axis indicates days post-injection and the y-axis indicates barcode level in urine. These plots show that late-shed kidney barcodes exhibit repeated detection across multiple urine timepoints, whereas late non-shed and random control barcodes are detected infrequently and typically at isolated timepoints.

We observed that individual late-shed kidney barcodes were repeatedly detected above the barcode-level detection threshold (weighted barcode count ≥10) across multiple urine timepoints in all four mice (**Fig. 3**). Many barcodes formed extended shedding trajectories spanning successive or nonconsecutive urine samples rather than isolated detection events. This sustained activity was consistent across animals, indicating that late shedding reflects stable, barcode-specific longitudinal behavior rather than dominance driven by a single transient peak.

In contrast, late non-shed kidney barcodes and randomly selected barcode controls were rarely detected in urine across time (**Fig. 3**). When late non-shed barcodes were detected, urine barcode levels were typically low and restricted to isolated timepoints. Although occasional late non-shed barcodes reached higher urine barcode levels, these events were sporadic and non-recurrent, failing to persist across subsequent urine collections. The longitudinal patterns of late non-shed barcodes closely resembled those of random controls (except in mouse “ML,” where late non-shed barcodes were still far less recurrent and sustained than late-shed barcodes), indicating that persistent urine shedding is specifically enriched among late-shed kidney barcodes rather than a generic consequence of barcode sampling or kidney residence.

### Divergence between late-shed and late non-shed kidney barcodes is not due to differences in input abundance

Stable, barcode-resolved differences in urine shedding raised the possibility that late-shed and late non-shed behaviors reflect differences already present in the viral inoculum. We tested this by examining the relative abundance of the top 50 kidney late-shed and top 50 late non-shed barcodes in the input virus stock. Because barcodes in the muPyV library are not evenly distributed in the input stock, as we have shown previously [31,32], we ranked barcodes by abundance in the inoculum and compared stock rank distributions for kidney late-shed and late non-shed barcodes across all four mice (**Fig. 4; Table S13-14**).

**Fig. 4.**
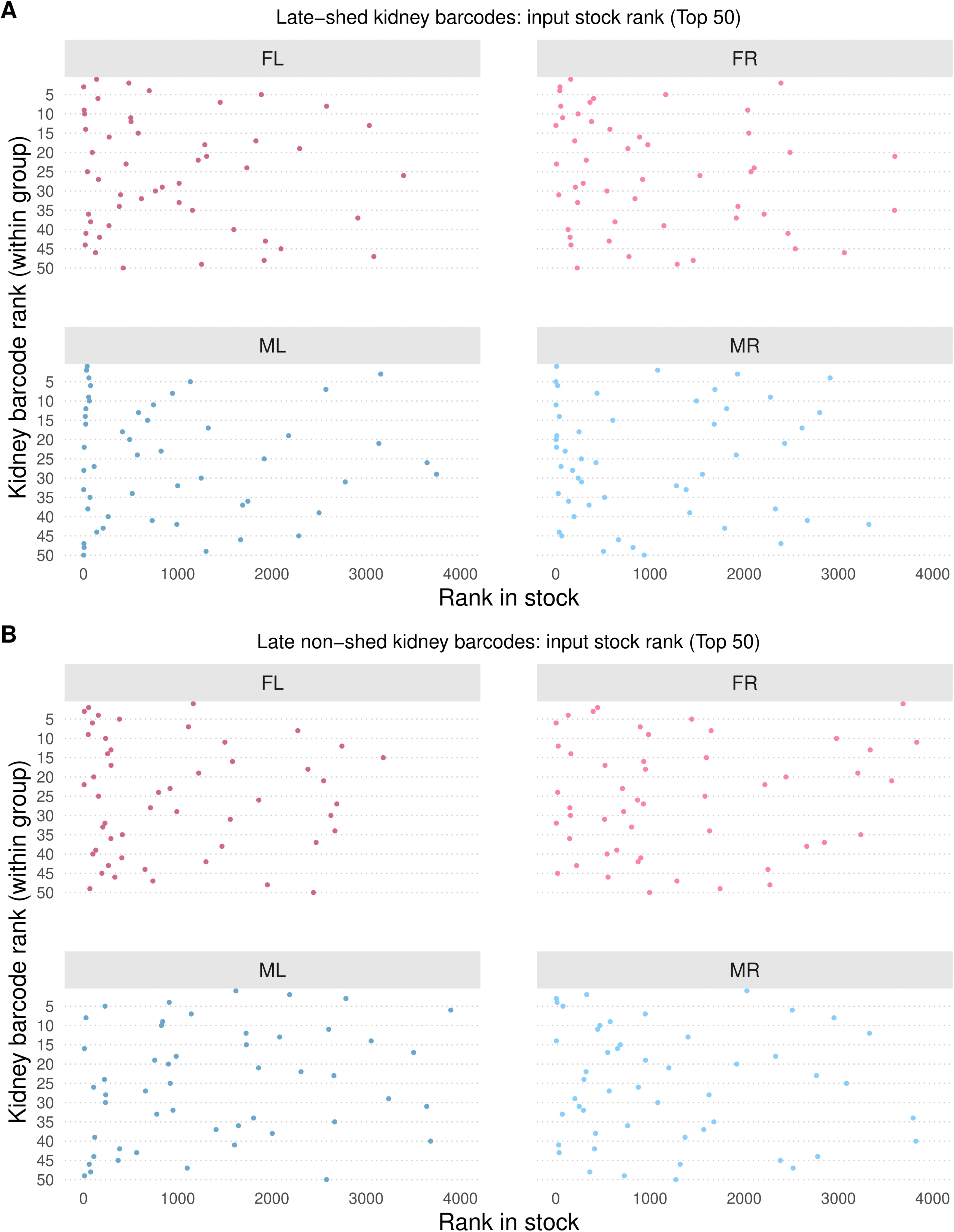
Input stock rank of late-shed and late non-shed kidney barcodes. Shown is the rank in the virus inoculum stock of kidney barcodes classified as late-shed or late non-shed, focusing on the top 50 barcodes per group in each mouse. In **(A)**, the input stock ranks of the top 50 late-shed kidney barcodes are shown for each mouse. In **(B)**, the input stock ranks of the top 50 late non-shed kidney barcodes are shown. A lower rank (toward the left side of the x-axis) indicates higher abundance in the virus stock, while the y-axis indicates the rank of each barcode within the kidney group. Note that late-shed and late non-shed kidney barcodes span broad and overlapping ranges of input stock rank across all four mice.

We found that kidney late-shed and late non-shed barcodes spanned broad and overlapping ranges of input stock ranks (**Fig. 4**). Across all four mice, the median input ranks of both groups fell within the top 25% of the inoculum, indicating that neither group was preferentially overrepresented at the time of infection (**Table S15-16**).

The absence of systematic stock-rank differences indicates that late shedding behavior emerges during infection rather than being predetermined by input abundance. If late-shed barcodes were preferentially selected simply due to higher input abundance, they would be expected to rank consistently much higher in the virus stock. Instead, late-shed and late non-shed kidney barcodes entered infection with comparable input abundance profiles, supporting a model in which early host-virus interactions establish divergent long-term shedding trajectories.

### Late-shed kidney barcodes reach substantially higher peak urine levels than late non-shed and random barcode controls

Sustained longitudinal shedding could arise either from repeated low-level activity or from occasional high-amplitude shedding events. We distinguished between these possibilities by comparing the maximum single-timepoint urine abundance reached by each barcode. For each mouse, we measured the highest urine barcode level observed across all timepoints for the top 50 late-shed kidney barcodes, the top 50 late non-shed kidney barcodes, and 50 randomly selected urine barcode controls, and compared these values across groups (**Fig. 5; Table S17**).

**Fig. 5.**
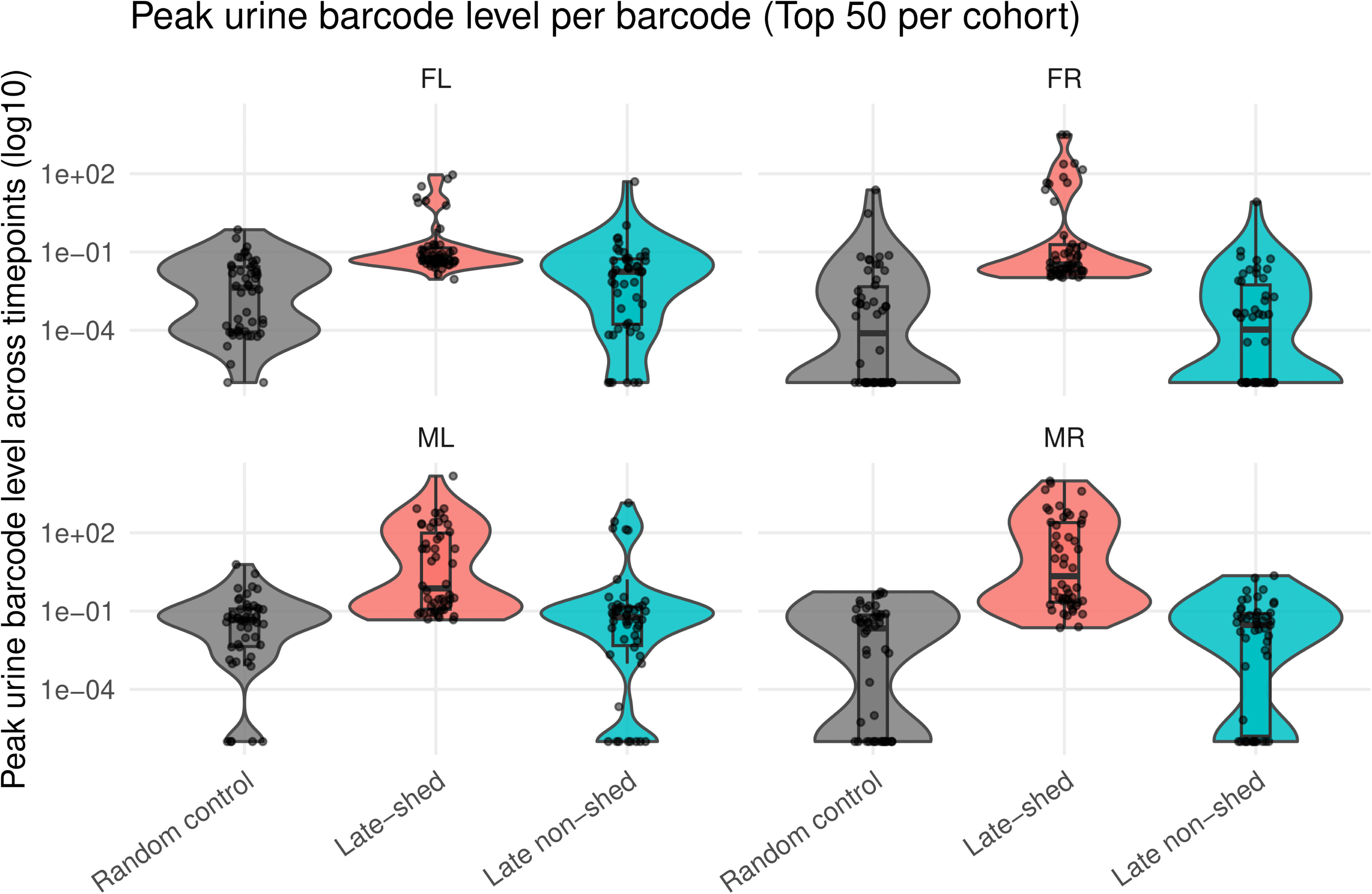
Peak urine levels of late-shed, late non-shed, and random barcode groups. Shown is the maximum urine barcode level reached by individual barcodes across all sampled urine timepoints for each mouse. For each mouse, peak urine levels were determined for the top 50 late-shed kidney barcodes, the top 50 late non-shed kidney barcodes, and a size-matched group of 50 randomly selected urine barcodes. Each point represents a single barcode, with values plotted on a log_₁₀_ scale. Violin plots depict the overall distribution of peak urine levels within each group, and box plots indicate the median and interquartile range. Note that late-shed kidney barcodes generally reach higher peak urine levels than late non-shed kidney barcodes and random barcode controls across all four mice.

We found that late-shed kidney barcodes consistently reached higher peak urine levels than both late non-shed kidney barcodes and random urine barcode controls across all four mice (**Fig. 5**). The distributions for late-shed kidney barcodes were shifted upward on a log scale, indicating a greater propensity to achieve high-amplitude urine detection at least once during infection.

In contrast, late non-shed kidney barcodes rarely reached high urine levels, even when considering their single largest detection event. Random urine barcode controls showed comparable distributions, with occasional low-level peaks but little evidence of high-amplitude shedding. These results indicate that high-amplitude urine shedding is a selective property of late-shed viruses, not a generic consequence of detection in urine.

### Top smoldering barcodes are selectively enriched among late-shed barcodes

Persistent, recurrent urine detection (“smoldering”) could reflect either stochastic barcode appearance or preferential activity of viral lineages associated with late shedding. We operationally defined smoldering based on detection frequency across urine timepoints. For each mouse and barcode, we calculated the fraction of urine timepoints in which that barcode was detected above the barcode-level detection threshold (weighted barcode count ≥10, where the weighted barcode count reflects the barcode’s read count after assignment to the closest matching stock barcode, with counts divided evenly among stock barcodes when a sample barcode matched multiple stock barcodes at the same minimum edit distance; see “Assigning sample barcodes to stock barcodes”). Using this metric, we defined the top smoldering barcodes as the 50 barcodes per mouse with the highest detection frequency and tested whether these barcodes were enriched among late-shed or late non-shed kidney barcodes compared with a size-matched random urine barcode baseline (**Fig. 6; Table S18**).

**Fig. 6.**
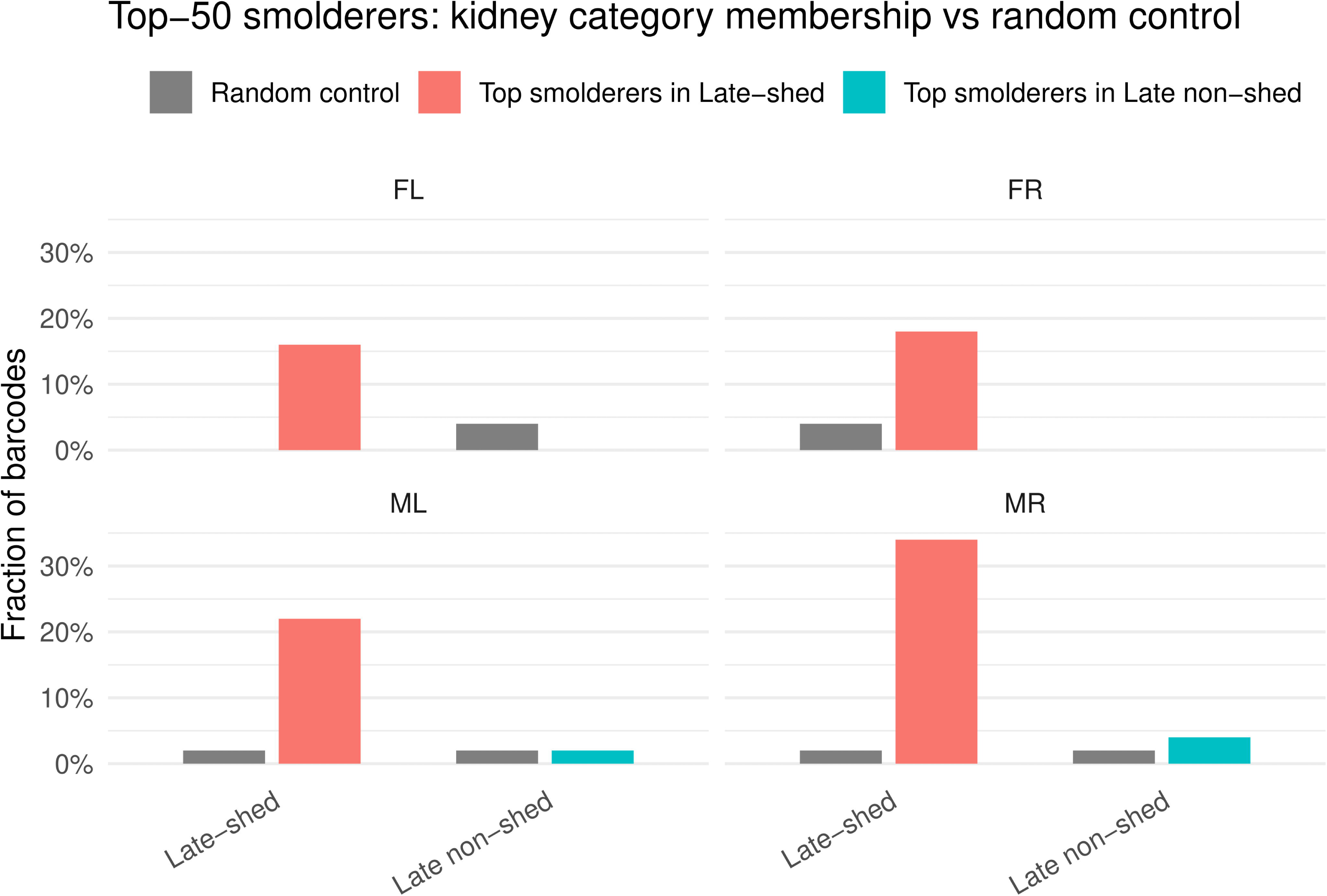
Distribution of top smoldering barcodes across kidney barcode categories. Smoldering was defined based on barcode detection frequency across urine timepoints, and for each mouse the top 50 barcodes with the highest fraction of urine timepoints detected were designated as top smoldering barcodes. Shown are the fractions of these top smoldering barcodes that fall into the late-shed kidney barcode group or the late non-shed kidney barcode group for each mouse compared with a size-matched random urine barcode control. Bars indicate the proportion of barcodes assigned to each kidney category, with random control distributions shown for reference. Note that, across all four mice, a larger fraction of top smoldering barcodes is found within the late-shed kidney barcode group than within the late non-shed group or the random control.

Across all four mice, we found that a substantially larger fraction of top smoldering barcodes belonged to the late-shed group than to the random urine control group (**Fig. 6**). In contrast, we observed little to no enrichment of top smoldering barcodes within the late non-shed group relative to random controls. This demonstrates that smoldering behavior is not randomly distributed across viral barcodes but is instead selectively enriched among kidney barcodes that ultimately contribute to late urinary shedding.

### Late-shed barcodes already exhibit elevated smoldering during early infection

While the analysis above tested whether barcodes with frequent detection across urine timepoints (“smoldering”) are enriched among late-shed kidney barcodes, it did not address whether this behavior is already apparent early after infection. To distinguish between early-established versus late-emerging shedding behavior, we next asked whether barcodes that ultimately dominate late urine already exhibit elevated detection during the early phase of infection. We partitioned each mouse’s urine time course by day, defining the late phase as the final quartile of urine sampling timepoints closest to the time of sacrifice (≥ day 74 for mice FL, FR, and ML; ≥ day 45 for mouse MR) and the early phase as all preceding.

For each mouse, we quantified early smoldering for three barcode groups: (i) the top 50 late smoldering barcodes, ranked by detection frequency above the barcode-level detection threshold (weighted barcode count ≥10; see “Assigning sample barcodes to stock barcodes”) during the late phase; (ii) the top 50 late high-shedding barcodes, ranked by total urine barcode level during the late phase; and (iii) a size-matched random urine barcode control group. For each barcode, we calculated the fraction of early-phase days in which it was detected above the detection threshold (**Fig. 7; Table S19**).

**Fig. 7.**
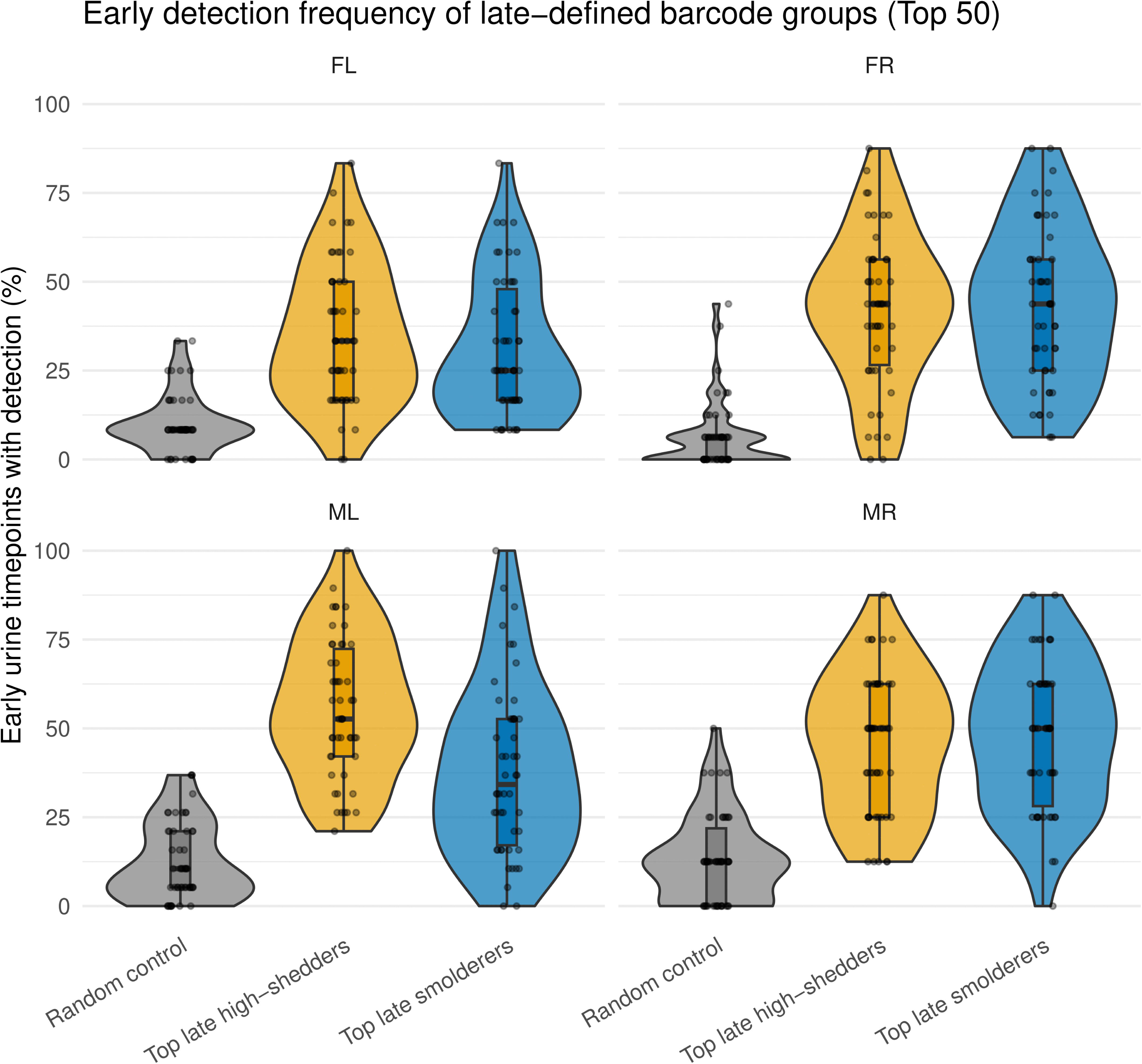
Early urine detection frequency of barcode groups defined by late-phase behavior. Urine time courses for each mouse were partitioned into an early phase and a late phase, with the late phase defined as urine samples collected on or after 75% of the total infection duration and the early phase defined as all preceding timepoints. Barcode groups were defined based on late-phase behavior, including the top 50 late smoldering barcodes (ranked by detection frequency during the late phase), the top 50 late high-shedding barcodes (ranked by total urine barcode level during the late phase), and a size-matched random urine barcode control group. Shown are the fractions of early-phase urine timepoints in which each barcode was detected, plotted for each group and mouse. Violin plots depict the distributions of early detection frequencies, with box plots indicating the median and interquartile range. Note that barcode groups defined by late-phase smoldering or high shedding show higher early detection frequencies than random barcode controls across all four mice.

We found that both late smoldering and late high-shedding barcodes exhibited substantially higher early detection frequencies than random urine barcodes across all four mice **(Fig. 7**). These results indicate that viral barcodes that ultimately dominate late urine, whether defined by persistent detection across urine timepoints or by high cumulative barcode level in urine, are already disproportionately detected in urine during early infection. Thus, by multiple criteria, we conclude that urinary shedding fate is established early during infection.

### The late non-shed kidney barcode population shows weak association with blood or other tissues

In our previous work, barcode relative abundance showed poor correlation across organs within the same mouse, indicating that productive viral populations are largely tissue-localized rather than freely exchanged [31]. In this study, we asked whether this principle extends to functionally distinct kidney barcode populations. To address this, we treated late-shed and late non-shed kidney barcodes as two pseudo-organ compartments and examined correlations in barcode relative abundance between these kidney-defined groups and other tissues within each mouse **(Fig. 8)**.

**Fig. 8.**
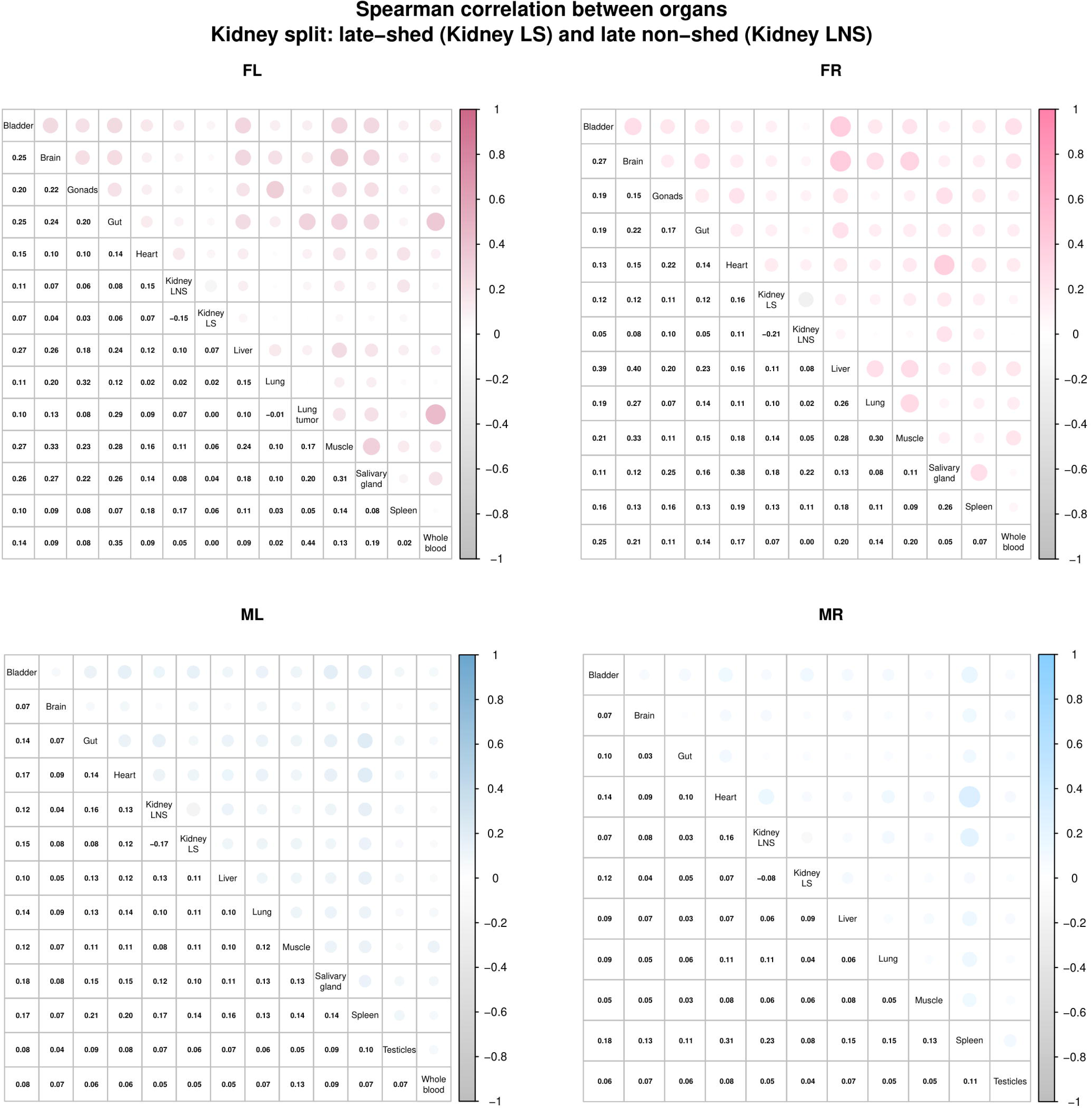
Low correlation of late-shed and late non-shed kidney barcode repertoires with other tissues. Shown is the Spearman correlation coefficient of relative barcode abundance between tissues within each mouse, with kidney barcodes partitioned into late-shed (Kidney LS) and late non-shed (Kidney LNS) groups. For each mouse, the top right portion of each matrix represents correlations by circle size and shading intensity between pairs of tissues, while the bottom left portion displays the corresponding numerical Spearman correlation coefficients. Correlations are shown between late-shed and late non-shed kidney barcode groups and other sampled tissues. Note that correlations between both late-shed and late non-shed kidney barcode groups and non-kidney tissues are generally low across all four mice.

Across all four mice, we observed weak correlations between barcode relative abundance in both late-shed and late non-shed kidney groups and abundance in non-kidney tissues (**Fig. 8**). Although some barcodes detected at high levels in one tissue were also detectable elsewhere, the identities of the most abundant barcodes differed between compartments, indicating that kidney-resident viral genomes segregate into functionally distinct populations whose abundance patterns are largely decoupled from those of other tissues.

Among the three mice for which whole-blood samples were available (FL, FR, and ML), we found little to no correlation between blood barcode abundance and late non-shed kidney barcodes, arguing against continuous hematogenous seeding of the silent kidney reservoir. Barcode relative abundance within both late-shed and late non-shed kidney populations also showed low correlation between animals, indicating that kidney barcode fates are independently established within each host (**Fig. S1**).

### Shedding and non-shedding fates arise independently of barcode sequence composition

Differences in late shedding behavior could arise from intrinsic features of the barcode sequence itself. We tested this possibility by examining multiple sequence features, including GC content (**Fig. S2; Table S20-21**), barcode length (**Fig. S3; Table S22**), and predicted host and viral miRNA seed matches (**Fig. S4; Table S23-24**).

For GC content and barcode length, we analyzed the top 50 kidney late-shed and top 50 kidney late non-shed barcodes per mouse. Across all mice, the GC-content distributions of late-shed and late non-shed barcodes closely overlapped with the global barcode distribution, indicating no enrichment for unusually high or low GC content **(Fig. S2)**. Barcode length distributions were also nearly identical between late-shed and late non-shed barcodes across all four mice, with no enrichment for specific barcode lengths **(Fig. S3)**.

Since we designed our barcodes to be incorporated into both early and late mRNAs in addition to tagging individual episomes, we determined whether host or viral miRNAs could be responsible for the different late-shed and late non-shed kidney virus populations. To determine whether miRNA-mediated targeting could bias shedding fate, we examined predicted seed matches for both host murine miRNAs and the muPyV-encoded miRNA across kidney-resident barcodes classified as late-shed or late non-shed. Specifically, miRNA seed matches were defined as perfect Watson-Crick complementarity to nucleotides 2-8 from the 5′ end of the mature miRNA. For host top-100 kidney abundant miRNAs [72], we quantified the fraction of kidney barcodes containing at least one predicted miRNA seed match, as well as the fraction of total kidney viral abundance attributable to miRNA-targeted barcodes, and compared these values between late-shed and late non-shed groups on a mouse-by-mouse basis. Across all four mice, both unweighted and abundance-weighted measures of host miRNA targeting were highly similar between late-shed and late non-shed barcode populations and fell within null distributions generated by length- and GC-matched resampling controls (see “Barcode sequence feature analyses”), indicating no enrichment or depletion beyond that expected from barcode sequence composition alone **(Fig. S4)**. In contrast, predicted targeting by the muPyV-encoded miRNA was extremely sparse, with only a very small number of kidney barcodes (0 to 4) exhibiting a predicted seed match across mice and groups **(Table S24)**.

These results indicate that barcode GC content, predicted miRNA targeting, and barcode length do not explain the divergence in shedding fate, supporting the conclusion that shedding and non-shedding behaviors arise from host- and infection-dependent processes rather than intrinsic properties of the barcode sequence.

## Discussion

PyV persistence is often framed at the level of the host, yet our barcode-resolved analyses show that within the kidney, individual viral genomes follow distinct and stable long-term trajectories. Across four mice, we consistently observe two kidney-resident barcode populations: a minority that repeatedly contributes to urine over long-term infection and a majority that persists in kidney without detectable contribution to late urine. This organization is not a trivial consequence of kidney barcode abundance, since late-shed and late non-shed barcodes span overlapping abundance ranges and many high-abundance kidney barcodes remain non-shedding. Nor is it explained by initial inoculum representation, as late-shed and late non-shed barcodes exhibit broad, overlapping stock-rank distributions. These findings indicate that long-term shedding behavior emerges during infection *in vivo*, rather than reflecting input abundance or kidney copy number.

A central observation of this study is that long-term urinary shedding reflects stable, barcode-resolved behavior rather than transient aggregate effects. Late-shed kidney barcodes were not simply detected at the final timepoints used for classification; instead, they tended to recur across multiple longitudinal urine samples, often forming extended shedding trajectories that were clearly distinguishable from the sparse, sporadic, near-random appearance of late non-shed kidney barcodes. Consistent with this, late-shed barcodes reached substantially higher peak urine levels than late non-shed or random urine controls. These results refine the population-level “smoldering plus bursts” framework from our previous study by demonstrating that persistent detection is not evenly distributed across kidney-resident genomes [31]. Rather, smoldering is selectively enriched among the kidney barcodes that also drive long-term urine output. The simplest interpretation is that a subset of kidney genomes occupies a high-shedding-competent trajectory that is established early and then maintained, producing both recurrent low-level detection and the capacity for high-amplitude episodes.

Support for early establishment comes from our finding that viral lineages dominating the late phase already exhibit elevated activity early after infection. Barcodes defined by frequent late-phase shedding (late smolderers) and those defined by late-phase total genome output (late high shedders) both showed increased detection frequencies early relative to random urine barcodes. This pattern is inconsistent with models in which the majority of high-output late shedding arises from late, stochastic activation of previously low-activity genomes [73–75]. Instead, it supports a framework in which early host-virus interactions set a long-term propensity to shed that becomes evident as persistent infection unfolds (**Fig. 9**). Importantly, this stability does not imply continuous productivity. Rather, it suggests that the probability of crossing a threshold into measurable urine output differs systematically among kidney-resident genomes.

**Fig. 9.**
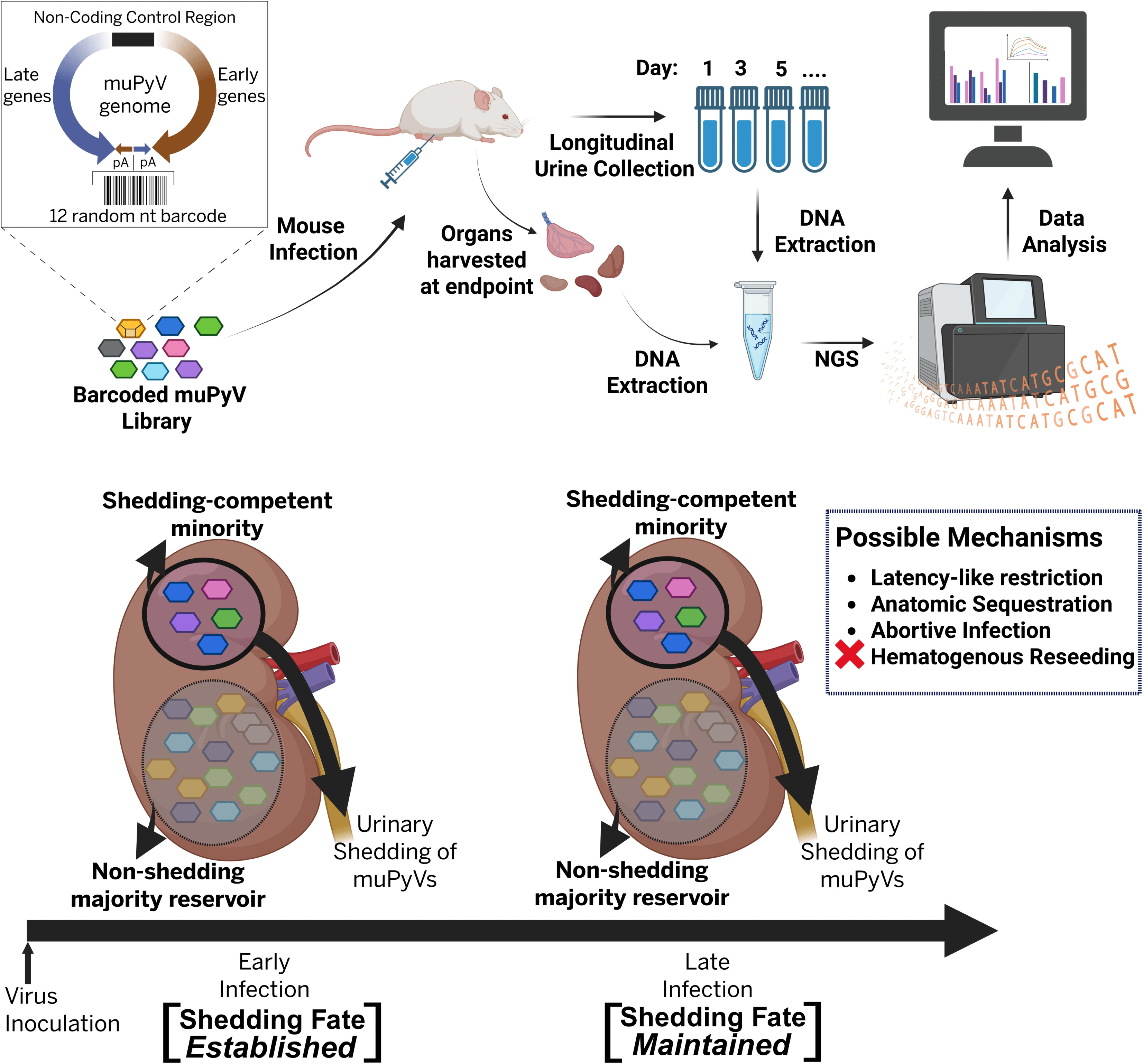
Conceptual model of lineage-resolved shedding fates during persistent muPyV infection. Shown is an overview of the experimental workflow and a conceptual model summarizing how kidney-resident viral barcodes diverge into long-term shedding-competent or non-shedding trajectories. The schematic outlines infection with a barcoded muPyV library, longitudinal urine collection, endpoint organ harvest, DNA extraction, and NGS-based barcode quantification. At the level of individual kidney-resident genomes, a minority of barcodes repeatedly contributes to urinary shedding, whereas a majority persists as a long-lived non-shedding reservoir. Early infection establishes barcode-specific shedding fates that are maintained through late infection. Potential mechanisms underlying this divergence include latency-like restriction, anatomical sequestration, and abortive infection, while hematogenous reseeding is disfavored by our data.

What biological mechanisms could generate a large, stable low/non-shedding reservoir alongside a smaller, repeatedly shedding-competent minority? A compelling explanation invokes latency-like or deeply restricted infection states gated by host cell state. PyV persistence has long been discussed in terms of both smoldering and latent/lytic–like infection programs, even though definitive evidence for true latency remains limited [76–80]. Recent work on BKPyV has shifted the field away from a purely virus-driven view of cell-cycle control toward a host-gated model, in which entry into S phase is required for robust LTAg expression and early transcription is shaped by Non-Coding Control Region (NCCR) responsiveness to host transcription factors including RB-E2F axis [37,38]. In this framework, the large late non-shed reservoir we observe is compatible with genomes persisting predominantly in host environments that rarely enter replication-competent states, whereas the late-shed minority resides in niches that intermittently permit early activation and can further reinforce replication competence through engagement of DNA-damage response pathways that prolong S phase [39,46].

Host immune signaling likely acts in parallel with cell-cycle gating to stabilize virus shedding patterns, serving as another determinant of viral fate. In particular, cell-type–specific antiviral programs, including type I interferon responses, may bias which kidney-resident environments remain deeply restricted versus permissive for activation [81]. Type I interferon signaling is especially intriguing in this regard, as it is known to stabilize latent or persistent states in other DNA viruses and has been shown to modulate PyV infection outcomes in a cell-type–dependent manner, suggesting a role in reinforcing restricted, non-shedding states rather than directly triggering reactivation [82–87]. Our data are consistent with a latency-like gating mechanism but do not establish true latency in the strict sense of reversible switching within the same cell. What we can conclude with confidence is that kidney-resident genomes behave as if partitioned into long-lived states with sharply different probabilities of producing urine-detectable output.

A second model, which our data more directly constrain, is hematogenous reseeding. Hematogenous dissemination has been proposed for human PyVs, particularly for JCPyV, where transient viremia and leukocyte-associated spread are thought to contribute to dissemination between peripheral reservoirs and the central nervous system [88–91]. If the non-shedding kidney reservoir were continuously replenished from blood, or if long-term urine contributors were repeatedly reseeded from systemic compartments, stronger coupling between barcode abundance in blood and kidney-defined populations would be expected. Instead, barcode compositions are largely tissue-local, likely due to robust IgG response [92], and blood shows little to no correlation with the late non-shed kidney population in the mice where blood was sampled. This argues against ongoing hematogenous input as a dominant driver of either the silent reservoir or the long-term shedders. Rather than being continually repopulated by incoming genomes, kidney-resident viral populations appear to be locally established and stably maintained within each host.

A third possibility is spatial or anatomical sequestration within the kidney [90,93–95]. In this model, many viral genomes are genuinely present but reside in regions poorly coupled to the urinary outflow, such that virions or viral nucleic acids rarely reach urine even if low-level replication occurs. Our data cannot exclude this explanation, as kidney abundance alone does not predict urinary contribution and the late non-shed population constitutes a major quantitative reservoir in most animals. Differences in PyV receptor distribution across kidney tissue types may also contribute to divergent infection outcomes within the kidney [96–98]. This idea is consistent with recent single-cell analyses of human BKPyV, which demonstrate ranked renal cell-type tropism rather than uniform infection across the kidney, indicating that different infected compartments may differ substantially in their likelihood of contributing to urine [48]. More broadly, it aligns with the idea that PyV shedding may reflect local tissue and cellular context, and that not all infected niches within a kidney-tropic infection are equally positioned to contribute to urine.

A fourth model is abortive infection at the level of individual genomes. Under this scenario, the late non-shed reservoir could be enriched for genomes that are genetically or functionally compromised, allowing persistence as DNA but rarely supporting productive replication. Our analyses argue against barcode-intrinsic explanations such as GC content, length, or predicted miRNA targeting, but they do not rule out defects elsewhere in the viral genome. Prior studies of PyVs have documented persistent viral populations containing rearranged or non-functional genomes [99–102]. Abortive-genome models therefore remain plausible, particularly if early immune pressure or cellular restriction favors survival of defective genomes. Any such model, however, must still account for the striking contrast between a small subset of barcodes that repeatedly achieves high peak urine levels and persistent detection and a much larger population that does not.

Our findings support a framework in which persistent kidney infection is not a homogeneous pool of equivalent genomes, but a structured ecosystem composed of at least two long-lived behavioral classes. The large late non-shed reservoir provides a plausible substrate for cryptic persistence, while the high-shedding-competent minority supplies sustained and episodic urinary output. This organization bridges classical descriptions of intermittent viruria with emerging host-gated models of PyV activity: persistent infection can appear intermittent at the host level because only a small, stable subset of genomes repeatedly reaches detectable shedding, while most remain abortive, spatially sequestered or silenced.

### Limitations of this study

This study is limited by sample size and by the indirect nature of urine as a readout of kidney biology. Although the four-mouse dataset provides internal replication of the core qualitative conclusions, larger cohorts will be required to assess sex effects, inter-host variability, and the generality of the observed proportions of shedding versus non-shedding reservoirs. This is particularly relevant given extensive evidence that sex hormones and endocrine-immune interactions can shape viral persistence and tissue-specific infection dynamics, including in muPyV models [103–105]. In addition, our classifications rely on urine sampling density and on defining “late shedding” using final timepoints proximal to harvest; more frequent urine sampling and time-resolved kidney measurements would improve sensitivity to rare events and better distinguish persistent low-level output from sporadic bursts. Finally, our barcode sequencing captures lineage identity and abundance but does not reveal full genome sequence, infected cell type, kidney microanatomical location, host cell state, or viral genome integrity. As a result, we cannot definitively discriminate among latency-like restriction, spatial sequestration, and abortive-genome models, which may coexist. Future studies linking barcode identity to cell state and spatial location, together with full-genome sequencing of shedding-competent and non-shedding reservoirs, will be essential for identifying the host and viral determinants that place a genome on a long-term shedding trajectory.

## Supplementary Figures

**Fig. S1. Low correlation of late-shed and late non-shed kidney barcode repertoires between mice.** Shown is the Spearman correlation of relative barcode abundance between mice for kidney barcodes classified as late-shed or late non-shed. For each group, pairwise correlations were calculated between animals using relative barcode abundance in kidney tissue. Circle size and shading intensity indicate the magnitude of the Spearman correlation coefficient, with numerical values shown in the corresponding matrix cells. Note that correlations between mice are low for both late-shed and late non-shed kidney barcode groups.

**Fig. S2. GC content of late-shed and late non-shed kidney barcodes.** Shown are the GC-content distributions of kidney barcodes classified as late-shed or late non-shed, focusing on the top 50 barcodes per group for each mouse. GC content is plotted as the percentage of guanine and cytosine nucleotides within each barcode. Box plots indicate the median and interquartile range, with individual points representing single barcodes. For reference, the GC-content distribution of all tissue barcodes is shown in gray. Note that GC-content distributions of late-shed and late non-shed kidney barcodes overlap with each other and with the global barcode distribution across all four mice.

**Fig. S3. Barcode length distributions of late-shed and late non-shed kidney barcodes.** Shown are the length distributions of kidney barcodes classified as late-shed or late non-shed, focusing on the top 50 barcodes per group for each mouse. Barcodes are grouped by nucleotide length, and bar heights indicate the number of barcodes of each length within each group. Note that barcode length distributions are similar between late-shed and late non-shed kidney barcodes across all four mice, with no enrichment for specific barcode lengths.

**Fig. S4. Host miRNA seed targeting in late-shed and late non-shed kidney barcode groups is consistent with the null distribution. (A)** Shown is the fraction of kidney-resident barcodes containing at least one seed (nt 2-8) match to the top 100 kidney-abundant host miRNAs, calculated separately for late-shed and late non-shed barcode groups for each mouse. Points indicate observed values for individual animals, with lines connecting late-shed and late non-shed groups within the same mouse. Violin plots show the null distribution generated by 2,000 resampling iterations in which barcodes were randomly reassigned within each mouse while preserving barcode length and GC-content distributions. **(B)** Same analysis as in (A), but weighted by kidney barcode abundance (barcode level), showing the fraction of total kidney viral abundance attributable to top 100 kidney-abundant host miRNA seed-targeted barcodes. In both unweighted and weighted analyses, values for late-shed and late non-shed groups overlap with each other and fall within the corresponding null distributions.

## Supplementary Tables

**Table S1.** Stock barcode abundance table showing barcodes retained above the 99th-percentile cutoff used for downstream analyses.

**Table S2.** Barcode-resolved urine sequencing data across all animals and timepoints, including normalized barcode abundance per sample.

**Table S3.** Barcode-resolved tissue sequencing data across all harvested organs, including normalized barcode abundance per tissue.

**Table S4.** Counts of kidney barcodes stratified by late-shed and late non-shed classification for each animal (Fig. 1A).

**Table S5.** Total kidney barcode abundance summarized by late-shed and late non-shed status across animals (Fig. 1B).

**Table S6.** Kidney barcode abundance table annotated with late-shed versus late non-shed classification (Fig. 1C).

**Table S7.** Data underlying donut plots showing urine barcode composition with highlighted top-50 late-shed kidney barcodes (Fig. 2A).

**Table S8.** Data underlying donut plots showing urine barcode composition with highlighted top-50 late non-shed kidney barcodes (Fig. 2B).

**Table S9.** Summed urine abundance of top-50 late-shed kidney barcodes over time for each animal (Fig. 2C).

**Table S10.** Summed urine abundance of top-50 late non-shed kidney barcodes over time for each animal (Fig. 2D).

**Table S11.** Lists of barcode sets used for clone-resolved urine trajectory analyses, including late-shed, late non-shed, and random control groups (Fig. 3).

**Table S12.** Time-resolved urine barcode trajectories for selected barcode sets used in Fig. 3.

**Table S13.** Stock-rank–annotated late-shed kidney barcodes, including per-barcode stock abundance ranks (Fig. 4).

**Table S14.** Stock-rank–annotated late non-shed kidney barcodes, including per-barcode stock abundance ranks (Fig. 4).

**Table S15.** Median stock abundance ranks of late-shed kidney barcodes calculated per animal.

**Table S16.** Median stock abundance ranks of late non-shed kidney barcodes calculated per animal.

**Table S17.** Peak urine shedding metrics for each barcode cohort across animals (Fig. 5).

**Table S18.** Enrichment statistics comparing top smoldering barcodes to random controls across kidney barcode categories (Fig. 6).

**Table S19.** Early-phase detection frequencies of late-smoldering, late high-shedding, and random barcode sets (Fig. 7).

**Table S20.** GC-content distribution of top-50 late-shed kidney barcodes compared to all tissue barcodes (Fig. S2A).

**Table S21.** GC-content distribution of top-50 late non-shed kidney barcodes compared to all tissue barcodes (Fig. S2B).

**Table S22.** Length distribution of top-ranked late-shed and late non-shed kidney barcodes (Fig. S3).

**Table S23.** Host (murine) miRNA seed matches (top 100 kidney-abundant miRNAs) in kidney barcodes classified as late-shed or late non-shed across mice (Fig. S4A).

**Table S24.** muPyV miRNA seed matches in kidney barcodes classified as late-shed or late non-shed across mice (Fig. S4B).

## Acknowledgments

We thank Sylvain Blois and Benni Goetz for contributing to the previously published raw data re-analyzed in this current study. We thank past and present members of the Sullivan Lab for assistance with mouse maintenance, urine collection, helpful discussions, guidance on data visualization, technical advice, and careful reading of the manuscript.

## Institutional Review Board Statement

All animal procedures were conducted in accordance with protocols approved by The University of Texas at Austin Institutional Animal Care and Use Committee (AUP-2022–00305) and were subject to veterinary approval.

## Data Availability Statement

All data underlying this study are provided within the figures, supplementary materials, and publicly accessible repositories. The raw sequencing data generated used in this study are available through the NCBI Sequence Read Archive (SRA) under BioProject accession number PRJNA791340. All custom analysis code and scripts has been made publicly available on GitHub at https://github.com/ChrisSullivanLab/Shedding-dynamics-of-a-DNA-virus within the *divergent.shedding.fate* directory. In addition, confirmatory documentation of independent validation of the major findings of this study (Figures 1A, 2, 3, 6, 7, 8, and S1), along with additional analyses performed by one of the authors (K.W.N.), have been made publicly available in the same directory of the GitHub repository.

## Author Contributions

**Conceptualization**: A.M. and C.S.S.; **Methodology**: A.M.; **Software**: A.M.; **Validation**: A.M. and K.W.N.; **Formal Analysis**: A.M., K.W.N., and C.S.S.; **Investigation**: A.M. and C.S.S.; **Resources**: C.S.S.; **Data Curation**: A.M.; **Writing – Original Draft Preparation**: A.M.; **Writing – Review & Editing**: A.M., K.W.N., and C.S.S.; **Visualization**: A.M. and K.W.N.; **Supervision**: C.S.S.; **Project Administration**: C.S.S.; **Funding Acquisition**: C.S.S.

## References

1. White, M.K.; Khalili, K. Polyomaviruses and Human Cancer: Molecular Mechanisms Underlying Patterns of Tumorigenesis. Virology 2004, 324, 1–16, doi:10.1016/j.virol.2004.03.025.

2. DiMaio, D., Small Size, Big Impact: How Studies of Small DNA Tumour Viruses Revolutionized Biology. Philos Trans R Soc Lond B Biol Sci 2019, 374, 20180300, doi:10.1098/rstb.2018.0300.

3. Pipas, J.M. DNA Tumor Viruses and Their Contributions to Molecular Biology. Journal of Virology 2019, 93, 10.1128/jvi.01524-18, doi:10.1128/jvi.01524-18.

4. Prado, J.C.M.; Monezi, T.A.; Amorim, A.T.; Lino, V.; Paladino, A.; Boccardo, E. Human Polyomaviruses and Cancer: An Overview. Clinics 2018, 73, e558s, 10.6061/clinics/2018/e558s.

5. Fanning, E.; Zhao, X.; Jiang, X. Polyomavirus Life Cycle. In DNA Tumor Viruses; Damania, B., Pipas, J.M., Eds.; Springer US: New York, NY, 2009; pp. 1–24 ISBN 978-0-387-68945-6.

6. Chang, Y.; Moore, P.S. Merkel Cell Carcinoma: A Virus-Induced Human Cancer. Annual Review of Pathology: Mechanisms of Disease 2012, 7, 123–144, doi:10.1146/annurev-pathol-011110-130227.

7. Arora, R.; Chang, Y.; Moore, P.S. MCV and Merkel Cell Carcinoma: A Molecular Success Story. Current Opinion in Virology 2012, 2, 489–498, doi:10.1016/j.coviro.2012.05.007.

8. Kamminga, S.; van der Meijden, E.; de Brouwer, C.; Feltkamp, M.; Zaaijer, H. Prevalence of DNA of Fourteen Human Polyomaviruses Determined in Blood Donors. Transfusion 2019, 59, 3689–3697, doi:10.1111/trf.15557.

9. Shen, C.-L.; Wu, B.-S.; Lien, T.-J.; Yang, A.-H.; Yang, C.-Y. BK Polyomavirus Nephropathy in Kidney Transplantation: Balancing Rejection and Infection. Viruses 2021, 13, 487, doi:10.3390/v13030487.

10. Alonso, M.; Villanego, F.; Orellana, C.; Vigara, L.A.; Montiel, N.; Aguilera, A.; Amaro, J.M.; Garcia, T.; Mazuecos, A. Impact of BK Polyomavirus Plasma Viral Load in Kidney Transplant Outcomes. Transplantation Proceedings 2022, 54, 2457–2461, doi:10.1016/j.transproceed.2022.10.012.

11. Rocchi, A.; Sariyer, I.K.; Berger, J.R. Revisiting JC Virus and Progressive Multifocal Leukoencephalopathy. J. Neurovirol. 2023, 29, 524–537, doi:10.1007/s13365-023-01164-w.

12. Lukacher, A.S.; Yuan, W.; O’Hara, B.A.; Garabian, K.; Haley, S.A.; Atwood, W.J. JC Polyomavirus Neuroinvasion across the Blood-Brain Barrier: Current Understanding and Emerging Perspectives. Tumour Virus Research 2025, 20, 200331, doi:10.1016/j.tvr.2025.200331.

13. Haley, S.A.; Atwood, W.J. Progressive Multifocal Leukoencephalopathy: Endemic Viruses and Lethal Brain Disease. Annual Review of Virology 2017, 4, 349–367, doi:10.1146/annurev-virology-101416-041439.

14. Hatton, G.H.; James, S.R.; Mason, A.S.; Gawne, R.T.; Vogel, H.; Hogg, K.; Boukani, P.; Swinscoe, G.; Khan, A.; Welberry Smith, M.;, et al. Virus-Induced APOBEC3 Transmutagenesis in Bladder Cancer Initiation. Science Advances 2025, 11, eaea6124, doi:10.1126/sciadv.aea6124.

15. Starrett, G.J.; Buck, C.B. BK Polyomavirus Is a Cause of Bladder Cancer. Curr Opin Virol 2019, 39, 8–15, doi:10.1016/j.coviro.2019.06.009.

16. Wendzicki, J.A.; Moore, P.S.; Chang, Y. Large T and Small T Antigens of Merkel Cell Polyomavirus. Current Opinion in Virology 2015, 11, 38–43, doi:10.1016/j.coviro.2015.01.009.

17. Richards, K.F.; Guastafierro, A.; Shuda, M.; Toptan, T.; Moore, P.S.; Chang, Y. Merkel Cell Polyomavirus T Antigens Promote Cell Proliferation and Inflammatory Cytokine Gene Expression. Journal of General Virology 2015, 96, 3532–3544, doi:10.1099/jgv.0.000287.

18. Harms, P.W.; Harms, K.L.; Moore, P.S.; DeCaprio, J.A.; Nghiem, P.; Wong, M.K.K.; Brownell, I. The Biology and Treatment of Merkel Cell Carcinoma: Current Understanding and Research Priorities. Nat Rev Clin Oncol 2018, 15, 763–776, doi:10.1038/s41571-018-0103-2.

19. Moore, P.S.; Chang, Y. Common Commensal Cancer Viruses. PLOS Pathogens 2017, 13, e1006078, doi:10.1371/journal.ppat.1006078.

20. Starrett, G.J.; Buck, C.B. The Case for BK Polyomavirus as a Cause of Bladder Cancer. Current Opinion in Virology 2019, 39, 8–15, doi:10.1016/j.coviro.2019.06.009.

21. Nickeleit, V.; Davis, V.G.; Thompson, B.; Singh, H.K. The Urinary Polyomavirus-Haufen Test: A Highly Predictive Non-Invasive Biomarker to Distinguish “Presumptive” from “Definitive” Polyomavirus Nephropathy: How to Use It—When to Use It—How Does It Compare to PCR Based Assays? Viruses 2021, 13, 135, doi:10.3390/v13010135.

22. Drachenberg, C.B.; Hirsch, H.H.; Papadimitriou, J.C.; Gosert, R.; Wali, R.K.; Munivenkatappa, R.; Nogueira, J.; Cangro, C.B.; Haririan, A.; Mendley, S.;, et al. Polyomavirus BK Versus JC Replication and Nephropathy in Renal Transplant Recipients: A Prospective Evaluation. Transplantation 2007, 84, 323, doi:10.1097/01.tp.0000269706.59977.a5.

23. Costa, C.; Cavallo, R. Polyomavirus-Associated Nephropathy. World J Transplant 2012, 2, 84–94, doi:10.5500/wjt.v2.i6.84.

24. Pipas, J.M.; Sullivan, C.S. mGem: Deciphering How Polyomaviruses Coexist with Their Hosts for a Lifetime. mBio 2026, 0, e02311–25, doi:10.1128/mbio.02311-25.

25. Urbano, P.R.P.; Oliveira, R.R.; Romano, C.M.; Pannuti, C.S.; Fink, M.C.D. da S. Occurrence, Genotypic Characterization, and Patterns of Shedding of Human Polyomavirus JCPyV and BKPyV in Urine Samples of Healthy Individuals in São Paulo, Brazil. Journal of Medical Virology 2016, 88, 153–158, doi:10.1002/jmv.24318.

26. Zhong, S.; Zheng, H.-Y.; Suzuki, M.; Chen, Q.; Ikegaya, H.; Aoki, N.; Usuku, S.; Kobayashi, N.; Nukuzuma, S.; Yasuda, Y.;, et al. Age-Related Urinary Excretion of BK Polyomavirus by Nonimmunocompromised Individuals. Journal of Clinical Microbiology 2007, 45, 193–198, doi:10.1128/jcm.01645-06.

27. Kling, C.L.; Wright, A.T.; Katz, S.E.; McClure, G.B.; Gardner, J.S.; Williams, J.T.; Meinerz, N.M.; Garcea, R.L.; Vanchiere, J.A. Dynamics of Urinary Polyomavirus Shedding in Healthy Adult Women. Journal of Medical Virology 2012, 84, 1459–1463, doi:10.1002/jmv.23319.

28. Burke, J.M.; Bass, C.R.; Kincaid, R.P.; Ulug, E.T.; Sullivan, C.S. The Murine Polyomavirus MicroRNA Locus Is Required To Promote Viruria during the Acute Phase of Infection. Journal of Virology 2018, 92, 10.1128/jvi.02131-17, doi:10.1128/jvi.02131-17.

29. Alexander, K.M.; Ayers, K.N.; Lukacher, A.E. Live Long and Persist: Polyomavirus Immune Evasion in the Brain and Kidney. Future Virology 2025, 20, 313–321, doi:10.1080/17460794.2025.2530832.

30. Barth, H.; Solis, M.; Kack-Kack, W.; Soulier, E.; Velay, A.; Fafi-Kremer, S. In Vitro and In Vivo Models for the Study of Human Polyomavirus Infection. Viruses 2016, 8, 292, doi:10.3390/v8100292.

31. Blois, S.; Goetz, B.M.; Mojumder, A.; Sullivan, C.S. Shedding Dynamics of a DNA Virus Population during Acute and Long-Term Persistent Infection. PLOS Pathogens 2025, 21, e1013083, doi:10.1371/journal.ppat.1013083.

32. Blois, S.; Goetz, B.M.; Bull, J.J.; Sullivan, C.S. Interpreting and De-Noising Genetically Engineered Barcodes in a DNA Virus. PLOS Computational Biology 2022, 18, e1010131, doi:10.1371/journal.pcbi.1010131.

33. Swanson, P.A.; Lukacher, A.E.; Szomolanyi-Tsuda, E. Immunity to Polyomavirus Infection: The Polyoma Virus-Mouse Model. Semin Cancer Biol 2009, 19, 244–251, doi:10.1016/j.semcancer.2009.02.003.

34. Hopcraft, S.E.; Damania, B. Tumour Viruses and Innate Immunity. Philos Trans R Soc Lond B Biol Sci 2017, 372, 20160267, doi:10.1098/rstb.2016.0267.

35. Lukacher, A.E. Pathogen-Host Standoff. Immunol Res 2004, 29, 139–150, doi:10.1385/IR:29:1-3:139.

36. Wilson, J.J.; Lin, E.; Pack, C.D.; Frost, E.L.; Hadley, A.; Swimm, A.I.; Wang, J.; Dong, Y.; Breeden, C.P.; Kalman, D.;, et al. Gamma Interferon Controls Mouse Polyomavirus Infection In Vivo. Journal of Virology 2011, 85, 10126–10134, doi:10.1128/jvi.00761-11.

37. Salisbury, N.J.H.; Patil-Amonkar, S.; Roman, A.; Galloway, D.A. E2F1-3 Activate Merkel Cell Polyomavirus Early Transcription and Replication 2025, 2025.11.15.688648.

38. Needham, J.M.; Perritt, S.E.; Thompson, S.R. Single-Cell Analysis Reveals Host S Phase Drives Large T Antigen Expression during BK Polyomavirus Infection. PLoS Pathog 2024, 20, e1012663, doi:10.1371/journal.ppat.1012663.

39. Justice, J.L.; Needham, J.M.; Thompson, S.R. BK Polyomavirus Activates the DNA Damage Response To Prolong S Phase. Journal of Virology 2019, 93, 10.1128/jvi.00130-19, doi:10.1128/jvi.00130-19.

40. An, P.; Robles, M.T.S.; Duray, A.M.; Cantalupo, P.G.; Pipas, J.M. Human Polyomavirus BKV Infection of Endothelial Cells Results in Interferon Pathway Induction and Persistence. PLOS Pathogens 2019, 15, e1007505, doi:10.1371/journal.ppat.1007505.

41. Caller, L.G.; Davies, C.T.R.; Antrobus, R.; Lehner, P.J.; Weekes, M.P.; Crump, C.M. Temporal Proteomic Analysis of BK Polyomavirus Infection Reveals Virus-Induced G2 Arrest and Highly Effective Evasion of Innate Immune Sensing. Journal of Virology 2019, 93, 10.1128/jvi.00595-19, doi:10.1128/jvi.00595-19.

42. Needham, J.M.; Greco, T.M.; Cristea, I.M.; Thompson, S.R. Ribosomal Protein S25 Promotes Cell Cycle Entry for a Productive BK Polyomavirus Infection. Philos Trans R Soc Lond B Biol Sci 2025, 380, 20230390, doi:10.1098/rstb.2023.0390.

43. Dyson, N.; Buchkovich, K.; Whyte, P.; Harlow, E. The Cellular 107K Protein That Binds to Adenovirus E1A Also Associates with the Large T Antigens of SV40 and JC Virus. Cell 1989, 58, 249–255, doi:10.1016/0092-8674(89)90839-8.

44. Sullivan, C.S.; Baker, A.E.; Pipas, J.M. Simian Virus 40 Infection Disrupts P130–E2F and P107–E2F Complexes but Does Not Perturb pRb–E2F Complexes. Virology 2004, 320, 218–228, doi:10.1016/j.virol.2003.10.035.

45. Tahseen, D.; Rady, P.L.; Tyring, S.K. Human Polyomavirus Modulation of the Host DNA Damage Response. Virus Genes 2020, 56, 128–135, doi:10.1007/s11262-020-01736-6.

46. Justice, J.L.; Needham, J.M.; Verhalen, B.; Jiang, M.; Thompson, S.R. BK Polyomavirus Requires the Mismatch Repair Pathway for DNA Damage Response Activation. Journal of Virology 2022, 96, e02028–21, doi:10.1128/jvi.02028-21.

47. Weissbach, F.H.; Follonier, O.M.; Schmid, S.; Leuzinger, K.; Schmid, M.; Hirsch, H.H. Single-Cell RNA-Sequencing of BK Polyomavirus Replication in Primary Human Renal Proximal Tubular Epithelial Cells Identifies Specific Transcriptome Signatures and a Novel Mitochondrial Stress Pattern. Journal of Virology 2024, 98, e01382–24, doi:10.1128/jvi.01382-24.

48. Yang, F.; Chen, X.; Zhang, H.; Zhao, G.-D.; Yang, H.; Qiu, J.; Meng, S.; Wu, P.; Tao, L.; Wang, Q.;, et al. Single-Cell Transcriptome Identifies the Renal Cell Type Tropism of Human BK Polyomavirus. International Journal of Molecular Sciences 2023, 24, 1330, doi:10.3390/ijms24021330.

49. An, P.; Cantalupo, P.G.; Zheng, W.; Sáenz-Robles, M.T.; Duray, A.M.; Weitz, D.; Pipas, J.M. Single-Cell Transcriptomics Reveals a Heterogeneous Cellular Response to BK Virus Infection. Journal of Virology 2021, 95, 10.1128/jvi.02237-20, doi:10.1128/jvi.02237-20.

50. Yang, F.; Zhang, H.; Chen, X.; Zhao, G.; Meng, S.; Li, Q.; Yin, D.; Huang, G. Integrating Single-Cell RNA-Seq and Bulk RNA-Seq Reveals Ischemic Injury Promoting Polyomavirus Replication by DNA Damage Response.

51. Zhao, G.-D.; Gao, R.; Hou, X.-T.; Zhang, H.; Chen, X.-T.; Luo, J.-Q.; Yang, H.-F.; Chen, T.; Shen, X.; Yang, S.-C.;, et al. Endoplasmic Reticulum Stress Mediates Renal Tubular Vacuolation in BK Polyomavirus-Associated Nephropathy. Front. Endocrinol. 2022, 13, doi:10.3389/fendo.2022.834187.

52. Fallatah, D.I.; Christmas, S.; Binshaya, A. Adaptive Immune Response and Evasion Strategies of BK Virus. Transplantation Reviews 2025, 39, 100959, doi:10.1016/j.trre.2025.100959.

53. Martin, M. Cutadapt Removes Adapter Sequences from High-Throughput Sequencing Reads. EMBnet.journal 2011, 17, 10–12, doi:10.14806/ej.17.1.200.

54. Wickham, H.; Averick, M.; Bryan, J.; Chang, W.; McGowan, L.D.; François, R.; Grolemund, G.; Hayes, A.; Henry, L.; Hester, J.;, et al. Welcome to the Tidyverse. Journal of Open Source Software 2019, 4, 1686, doi:10.21105/joss.01686.

55. Wilke, C. Ggridges: Ridgeline Plots in “Ggplot2.”

56. Zorita, E.; Cuscó, P.; Filion, G.J. Starcode: Sequence Clustering Based on All-Pairs Search. Bioinformatics 2015, 31, 1913–1919, doi:10.1093/bioinformatics/btv053.

57. Pedersen, T.L. Patchwork: The Composer of Plots 2025.

58. Wilke, C.O. Ggridges: Ridgeline Plots in “Ggplot2”; 2025;

59. Lex, A.; Gehlenborg, N.; Strobelt, H.; Vuillemot, R.; Pfister, H. UpSet: Visualization of Intersecting Sets. IEEE Transactions on Visualization and Computer Graphics 2014, 20, 1983–1992, doi:10.1109/TVCG.2014.2346248.

60. Kolde, R. Pheatmap: Pretty Heatmaps; 2025;

61. Wei, T.; Simko, V. R Package “Corrplot”: Visualization of a Correlation Matrix; 2024;

62. Zhu, H. kableExtra: Construct Complex Table with Kable and Pipe Syntax 2024.

63. Loo, M.P.J. van der The Stringdist Package for Approximate String Matching. The R Journal 2014, 6, 111–122.

64. Wickham, H.; Henry, L. Purrr: Functional Programming Tools; 2025;

65. Wickham, H. Reshaping Data with the Reshape Package. Journal of Statistical Software 2007, 21, 1–20.

66. Auguie, B. gridExtra: Miscellaneous Functions for “Grid” Graphics; 2017;

67. Ameres, S.L.; Martinez, J.; Schroeder, R. Molecular Basis for Target RNA Recognition and Cleavage by Human RISC. Cell 2007, 130, 101–112, doi:10.1016/j.cell.2007.04.037.

68. Gorski, S.A.; Vogel, J.; Doudna, J.A. RNA-Based Recognition and Targeting: Sowing the Seeds of Specificity. Nat Rev Mol Cell Biol 2017, 18, 215–228, doi:10.1038/nrm.2016.174.

69. John, B.; Enright, A.J.; Aravin, A.; Tuschl, T.; Sander, C.; Marks, D.S. Human MicroRNA Targets. PLOS Biology 2004, 2, e363, doi:10.1371/journal.pbio.0020363.

70 Kozomara, A.; Birgaoanu, M.; Griffiths-Jones, S. miRBase: From microRNA Sequences to Function. Nucleic Acids Res 2019, 47, D155–D162, doi:10.1093/nar/gky1141.

71. Burke, J.M.; Bass, C.R.; Kincaid, R.P.; Ulug, E.T.; Sullivan, C.S. The Murine Polyomavirus MicroRNA Locus Is Required To Promote Viruria during the Acute Phase of Infection. Journal of Virology 2018, doi:10.1128/jvi.02131-17.

72. Isakova, A.; Fehlmann, T.; Keller, A.; Quake, S.R. A Mouse Tissue Atlas of Small Noncoding RNA. Proceedings of the National Academy of Sciences 2020, 117, 25634–25645, doi:10.1073/pnas.2002277117.

73. Yao, W.; Hertel, L.; Wahl, L.M. Dynamics of Recurrent Viral Infection. Proc Biol Sci 2006, 273, 2193–2199, doi:10.1098/rspb.2006.3563.

74. Singh, A.; Weinberger, L.S. Stochastic Gene Expression as a Molecular Switch for Viral Latency. Current Opinion in Microbiology 2009, 12, 460–466, doi:10.1016/j.mib.2009.06.016.

75. Rong, L.; Perelson, A.S. Modeling Latently Infected Cell Activation: Viral and Latent Reservoir Persistence, and Viral Blips in HIV-Infected Patients on Potent Therapy. PLOS Computational Biology 2009, 5, e1000533, doi:10.1371/journal.pcbi.1000533.

76. Swanson, P.A.; Lukacher, A.E.; Szomolanyi-Tsuda, E. Immunity to Polyomavirus Infection: The Polyomavirus–Mouse Model. Seminars in Cancer Biology 2009, 19, 244–251, doi:10.1016/j.semcancer.2009.02.003.

77. Lauver, M.D.; Katz, Z.E.; Markus, H.; Derosia, N.M.; Jin, G.; Ayers, K.N.; Butic, A.B.; Bushey, K.; Abendroth, C.S.; Liu, D.J.;, et al. The CXCR6-CXCL16 Axis Mediates T Cell Control of Polyomavirus Infection in the Kidney. PLOS Pathogens 2025, 21, e1012969, doi:10.1371/journal.ppat.1012969.

78. Heritage, J.; Chesters, P.M.; McCance, D.J. The Persistence of Papovavirus BK DNA Sequences in Normal Human Renal Tissue. Journal of Medical Virology 1981, 8, 143–150, doi:10.1002/jmv.1890080208.

79. Kwun, H.J.; Chang, Y.; Moore, P.S. Protein-Mediated Viral Latency Is a Novel Mechanism for Merkel Cell Polyomavirus Persistence. Proceedings of the National Academy of Sciences 2017, 114, E4040–E4047, doi:10.1073/pnas.1703879114.

80. Monaco, M.C.; Atwood, W.J.; Gravell, M.; Tornatore, C.S.; Major, E.O. JC Virus Infection of Hematopoietic Progenitor Cells, Primary B Lymphocytes, and Tonsillar Stromal Cells: Implications for Viral Latency. J Virol 1996, 70, 7004–7012, doi:10.1128/JVI.70.10.7004-7012.1996.

81. Qin, Q.; Shwetank, null; Frost, E.L.; Maru, S.; Lukacher, A.E. Type I Interferons Regulate the Magnitude and Functionality of Mouse Polyomavirus-Specific CD8 T Cells in a Virus Strain-Dependent Manner. J Virol 2016, 90, 5187–5199, doi:10.1128/JVI.00199-16.

82. An, P.; Robles, M.T.S.; Duray, A.M.; Cantalupo, P.G.; Pipas, J.M. Human Polyomavirus BKV Infection of Endothelial Cells Results in Interferon Pathway Induction and Persistence. PLOS Pathogens 2019, 15, e1007505, doi:10.1371/journal.ppat.1007505.

83. Cuddy, S.R.; Cliffe, A.R. The Intersection of Innate Immune Pathways with the Latent Herpes Simplex Virus Genome. Journal of Virology 2023, 97, e01352–22, doi:10.1128/jvi.01352-22.

84. Ryabchenko, B.; Soldatova, I.; Šroller, V.; Forstová, J.; Huérfano, S. Immune Sensing of Mouse Polyomavirus DNA by P204 and cGAS DNA Sensors. The FEBS Journal 2021, 288, 5964–5985, doi:10.1111/febs.15962.

85. Qin, Q.; Shwetank; Frost, E.L.; Maru, S.; Lukacher, A.E. Type I Interferons Regulate the Magnitude and Functionality of Mouse Polyomavirus-Specific CD8 T Cells in a Virus Strain-Dependent Manner. Journal of Virology 2016, 90, 5187–5199, doi:10.1128/jvi.00199-16.

86. Wu, H.-H.; Li, Y.-J.; Weng, C.-H.; Hsu, H.-H.; Chang, M.-Y.; Yang, H.-Y.; Yang, C.-W.; Tian, Y.-C. Interferon-Alpha and MxA Inhibit BK Polyomavirus Replication by Interaction with Polyomavirus Large T Antigen. Biomedical Journal 2024, 47, 100682, doi:10.1016/j.bj.2023.100682.

87. Assetta, B.; De Cecco, M.; O’Hara, B.; Atwood, W.J. JC Polyomavirus Infection of Primary Human Renal Epithelial Cells Is Controlled by a Type I IFN-Induced Response. mBio 2016, *7*, 10.1128/mbio.00903-16, doi:10.1128/mbio.00903-16.

88. Wollebo, H.S.; White, M.K.; Gordon, J.; Berger, J.R.; Khalili, K. Persistence and Pathogenesis of the Neurotropic Polyomavirus JC. Annals of Neurology 2015, 77, 560–570, doi:10.1002/ana.24371.

89. Imperiale, M.J.; Jiang, M. Polyomavirus Persistence. Annual Review of Virology 2016, 3, 517–532, doi:10.1146/annurev-virology-110615-042226.

90. Doerries, K. Human Polyomavirus JC and BK Persistent Infection. In Polyomaviruses and Human Diseases; Ahsan, N., Ed.; Springer: New York, NY, 2006; pp. 102–116 ISBN 978-0-387-32957-4.

91. Van Loy, T.; Thys, K.; Ryschkewitsch, C.; Lagatie, O.; Monaco, M.C.; Major, E.O.; Tritsmans, L.; Stuyver, L.J. JC Virus Quasispecies Analysis Reveals a Complex Viral Population Underlying Progressive Multifocal Leukoencephalopathy and Supports Viral Dissemination via the Hematogenous Route. Journal of Virology 2014, 89, 1340–1347, doi:10.1128/jvi.02565-14.

92. Ayers, K.N.; Lauver, M.D.; Alexander, K.M.; Jin, G.; Paraiso, K.; Ochetto, A.; Garg, S.; Goetschius, D.J.; Hafenstein, S.L.; Wang, J.C.-Y.; et al. The CD4 T Cell-Independent IgG Response during Persistent Virus Infection Favors Emergence of Neutralization-Escape Variants 2024, 2024.12.22.629980.

93. Alcendor, D.J. BK Polyomavirus Virus Glomerular Tropism: Implications for Virus Reactivation from Latency and Amplification during Immunosuppression. Journal of Clinical Medicine 2019, 8, 1477, doi:10.3390/jcm8091477.

94. Hirsch, H.H.; Steiger, J. Polyomavirus BK. The Lancet Infectious Diseases 2003, 3, 611–623, doi:10.1016/S1473-3099(03)00770-9.

95. Lorentzen, E.M.; Henriksen, S.; Rinaldo, C.H. Modelling BK Polyomavirus Dissemination and Cytopathology Using Polarized Human Renal Tubule Epithelial Cells. PLOS Pathogens 2023, 19, e1011622, doi:10.1371/journal.ppat.1011622.

96. Haley, S.A.; O’Hara, B.A.; Nelson, C.D.S.; Brittingham, F.L.P.; Henriksen, K.J.; Stopa, E.G.; Atwood, W.J. Human Polyomavirus Receptor Distribution in Brain Parenchyma Contrasts with Receptor Distribution in Kidney and Choroid Plexus. Am J Pathol 2015, 185, 2246–2258, doi:10.1016/j.ajpath.2015.04.003.

97. O’Hara, S.D.; Garcea, R.L. Murine Polyomavirus Cell Surface Receptors Activate Distinct Signaling Pathways Required for Infection. mBio 2016, 7, e01836–16, doi:10.1128/mBio.01836-16.

98. O’Hara, S.D.; Stehle, T.; Garcea, R. Glycan Receptors of the *Polyomaviridae*: Structure, Function, and Pathogenesis. Current Opinion in Virology 2014, 7, 73–78, doi:10.1016/j.coviro.2014.05.004.

99. Kodama, M.; Kojima, K.; Kodama, T. Studies on L Cells Persistently Infected With Polyoma Virus: Relationship Between D Particle and Other Virus-Like Particles2. J Natl Cancer Inst 1968, 41, 565–579, doi:10.1093/jnci/41.2.565.

100. Olsen, G.-H.; Hirsch, H.H.; Rinaldo, C.H. Functional Analysis of Polyomavirus BK Non-Coding Control Region Quasispecies from Kidney Transplant Recipients. Journal of Medical Virology 2009, 81, 1959–1967, doi:10.1002/jmv.21605.

101. Imperiale, M.J.; Jiang, M. What DNA Viral Genomic Rearrangements Tell Us about Persistence. Journal of Virology 2015, 89, 1948–1950, doi:10.1128/jvi.01227-14.

102. Cubitt, C.L. Molecular Genetics of the BK Virus. Adv Exp Med Biol 2006, 577, 85–95, doi:10.1007/0-387-32957-9_6.

103. Klein, S.L. The Effects of Hormones on Sex Differences in Infection: From Genes to Behavior. Neuroscience & Biobehavioral Reviews 2000, 24, 627–638, doi:10.1016/S0149-7634(00)00027-0.

104. McCance, D.J.; Mims, C.A. Reactivation of Polyoma Virus in Kidneys of Persistently Infected Mice during Pregnancy. Infection and Immunity 1979, 25, 998–1002, doi:10.1128/iai.25.3.998-1002.1979.

105. Wirth, J.J.; Amalfitano, A.; Gross, R.; Oldstone, M.B.; Fluck, M.M. Organ- and Age-Specific Replication of Polyomavirus in Mice. Journal of Virology 1992, 66, 3278–3286, doi:10.1128/jvi.66.6.3278-3286.1992.

