## Supplementary material for "Divergent Fates of Kidney-Resident Polyomaviruses: Stable Shedding Versus Near-Silent Persistence": Supplementary Figures.pdf

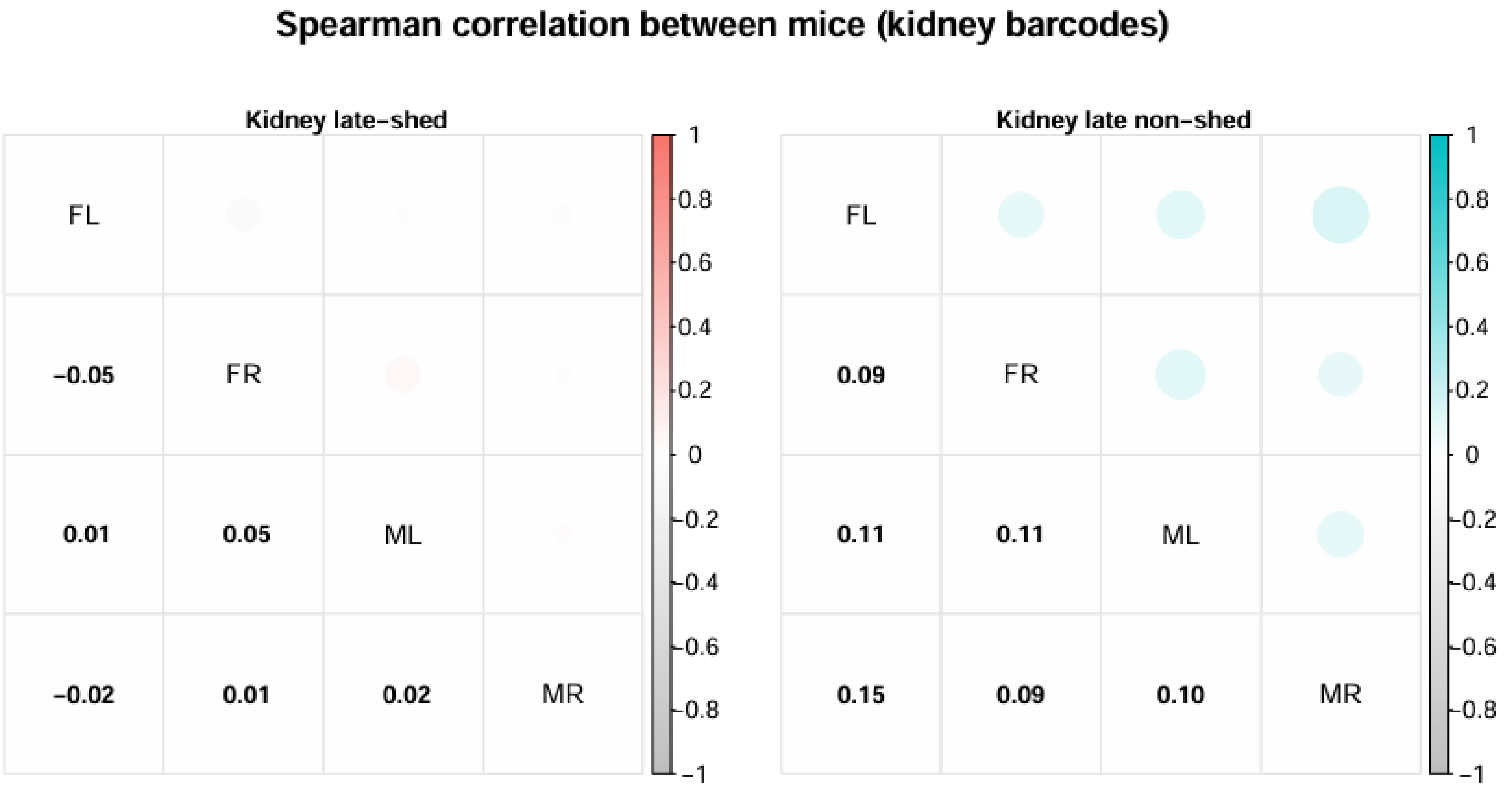


**Fig. S1. Low correlation of late-shed and late non-shed kidney barcode repertoires between mice.** Shown is the Spearman correlation of relative barcode abundance between mice for kidney barcodes classified as late-shed or late non-shed. For each group, pairwise correlations were calculated between animals using relative barcode abundance in kidney tissue. Circle size and shading intensity indicate the magnitude of the Spearman correlation coefficient, with numerical values shown in the corresponding matrix cells. Note that correlations between mice are low for both late-shed and late non-shed kidney barcode groups.

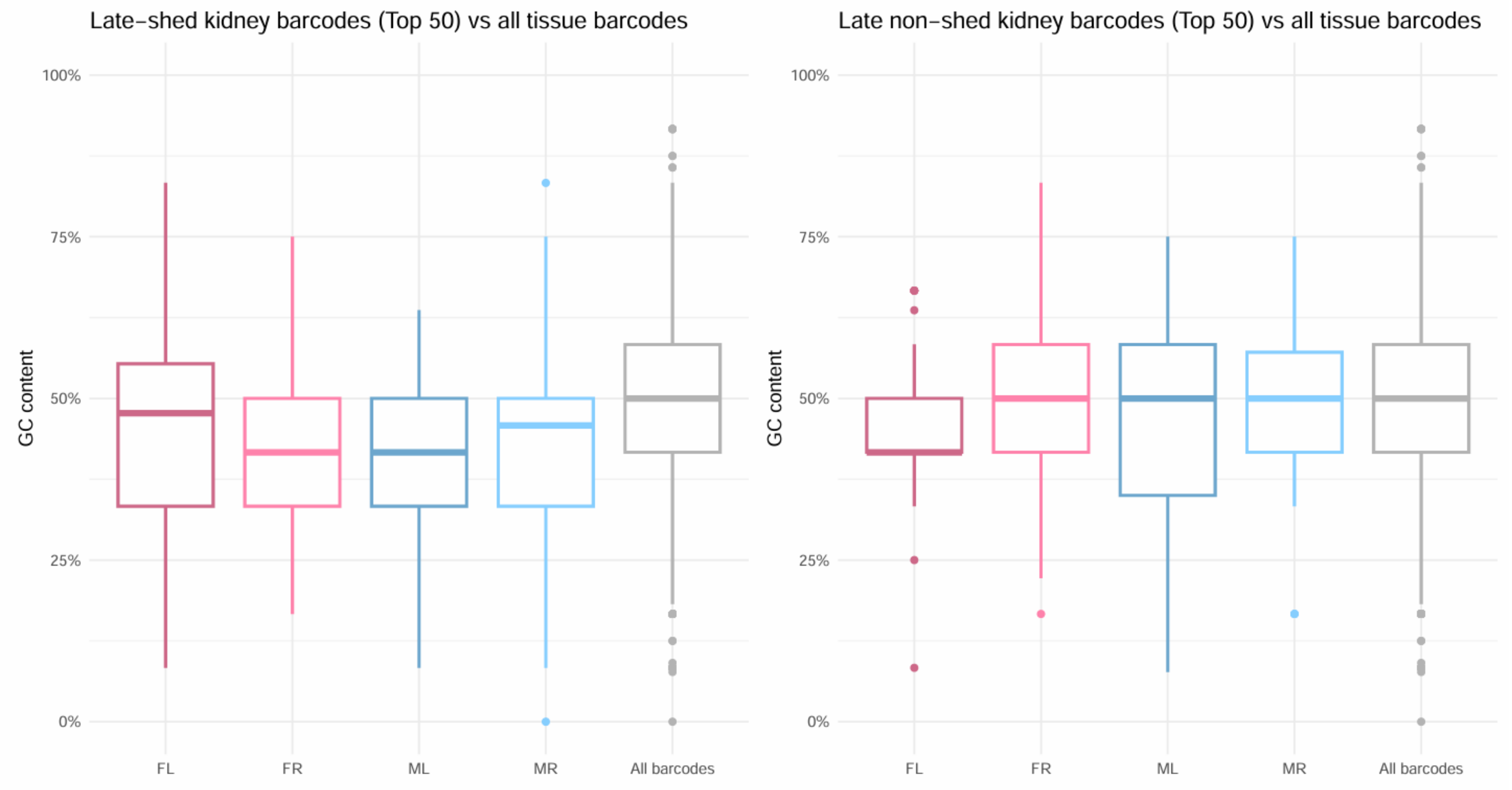


**Fig. S2. GC content of late-shed and late non-shed kidney barcodes.** Shown are the GC-content distributions of kidney barcodes classified as late-shed or late non-shed, focusing on the top 50 barcodes per group for each mouse. GC content is plotted as the percentage of guanine and cytosine nucleotides within each barcode. Box plots indicate the median and interquartile range, with individual points representing single barcodes. For reference, the GC-content distribution of all tissue barcodes is shown in gray. Note that GC-content distributions of late-shed and late non-shed kidney barcodes overlap with each other and with the global barcode distribution across all four mice.

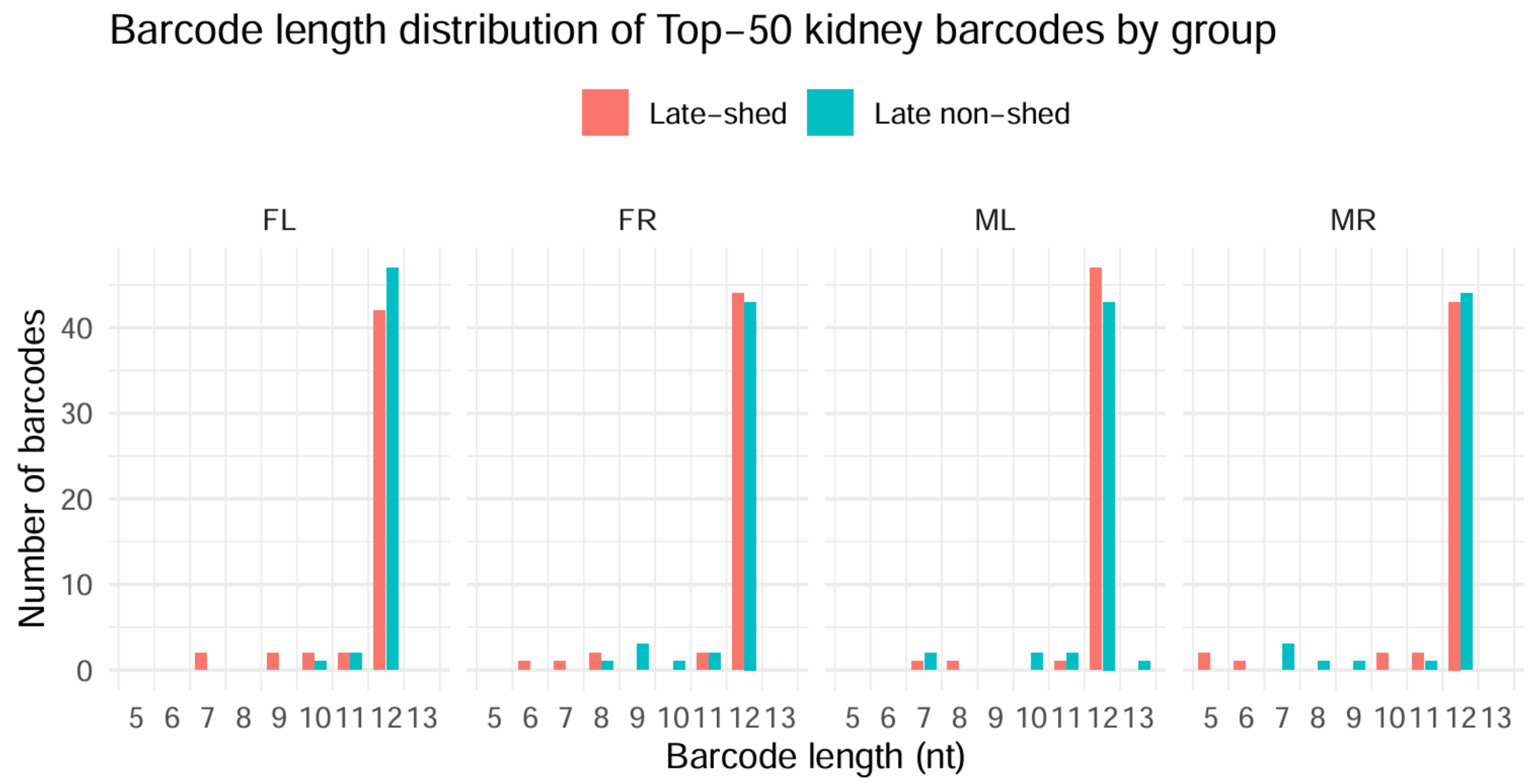


**Fig. S3. Barcode length distributions of late-shed and late non-shed kidney barcodes.** Shown are the length distributions of kidney barcodes classified as late-shed or late non-shed, focusing on the top 50 barcodes per group for each mouse. Barcodes are grouped by nucleotide length, and bar heights indicate the number of barcodes of each length within each group. Note that barcode length distributions are similar between late-shed and late non-shed kidney barcodes across all four mice, with no enrichment for specific barcode lengths.

**Fig. S4. Host miRNA seed targeting in late-shed and late non-shed kidney barcode groups is consistent with the null distribution. (A)** Shown is the fraction of kidney-resident barcodes containing at least one seed (nt 2-8) match to the top 100 kidney-abundant host miRNAs, calculated separately for late-shed and late non-shed barcode groups for each mouse. Points indicate observed values for individual animals, with lines connecting late-shed and late non-shed groups within the same mouse. Violin plots show the null distribution generated by 2,000 resampling iterations in which barcodes were randomly reassigned within each mouse while preserving barcode length and GC-content distributions. **(B)** Same analysis as in (A), but weighted by kidney barcode abundance (barcode level), showing the fraction of total kidney viral abundance attributable to top 100 kidney-abundant host miRNA seed-targeted barcodes. In both unweighted and weighted analyses, values for late-shed and late non-shed groups overlap with each other and fall within the corresponding null distributions.

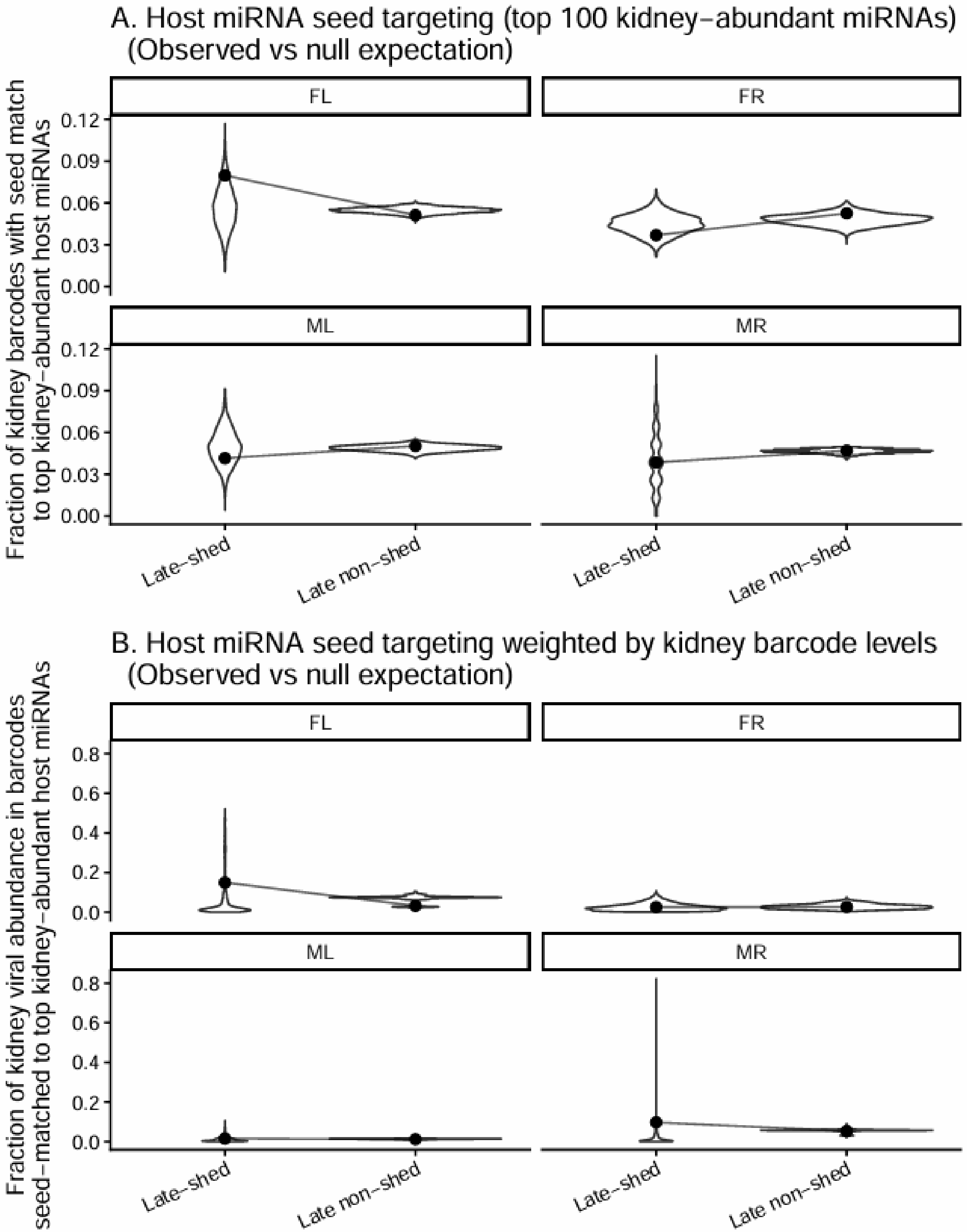
